# Transcriptome of the synganglion and characterization of the cys-loop ligand-gated ion channel gene family in the tick *Ixodes ricinus*

**DOI:** 10.1101/2021.12.20.473502

**Authors:** Claude Rispe, Caroline Hervet, Nathalie de la Cotte, Romain Daveu, Karine Labadie, Benjamin Noël, Jean-Marc Aury, Steeve Thany, Emiliane Taillebois, Alison Cartereau, Anaïs Le Mauff, Claude L. Charvet, Clément Auger, Cédric Neveu, Olivier Plantard

## Abstract

Ticks represent a major health issue for humans and domesticated animals. Assessing the expression patterns of the tick’s central nervous system, known as the synganglion, is an important step in understanding tick physiology and in managing tick-borne diseases. Neuron-specific genes like the cys-loop ligand-gated ion channels (cys-loop LGICs) are important pharmacological targets of acaricides. Here, we carried out the sequencing of transcriptomes of the *I. ricinus* synganglion, for adult ticks in different conditions (unfed males, unfed females, and partially-fed females). The *de novo* assembly of these transcriptomes allowed us to obtain a large collection of cys-loop LGICs sequences. A reference meta-transcriptome based on synganglion and whole body transcriptomes was then produced, showing high completeness and allowing differential expression analyses between synganglion and whole body. Many of the genes upregulated in the synganglion were related to biological processes or functions associated with neurotransmission and located in neurones or the synaptic membrane, including most of the cys-loop LGICs. As a first step of a functional study of cysLGICs, we cloned the predicted sequence of the resistance to dieldrin (RDL) subunit homologue, and functionally reconstituted the first GABA-gated receptor of *Ixodes ricinus* using a hetrologous expression approach. A phylogenetic study was performed for the nicotinic acetylcholine receptors (nAChRs) and for other cys-loop LGICs respectively, showing tick-specific expansions of some types of receptors (Histamine-gated, GluCls).

## Introduction

Like the vertebrate brain, the central nervous system (CNS) of arthropods performs essential functions, such as the perception of stimuli, the control of movement and the regulation of many physiological processes. Whether one compares the brain of vertebrates and the CNS of arthropods, or compares the CNS of different arthropod groups, important similarities but also differences are evident, and can be related to the specific biology and evolution of each group^1^. In ticks, a group of haematophagous arthropods belonging to the Chelicerata, the CNS is a specialized organ called a synganglion^2^, organized as a single mass of cells. As ticks are potential transmitters of many pathogens and parasites, a better knowledge of the function of this essential organ could help in understanding interactions between ticks and tick-borne-pathogens^3^, but also provide targets for chemical acaricides^4^. Indeed, to-date, most acaricides have targeted neural functions, but as with many insect pests, the continued use of these molecules has generated resistance, prompting the need to find new methods or targets^5^.

Knowledge of the neural biology of ticks is scarced compared to other invertebrates^2^, particularly with respect to gene expression changes in the synganglion during blood feeding or after mating^6.^. However progress has been made, notably through the sequencing of synganglion transcriptomes of several tick species. For example, a study of expressed-sequence tags of the *Rhipicephalus sanguineus* synganglion has identified different neural receptors^7^. Two companion studies of this study reported the sequencing of synganglion transcriptomes of the American dog tick, *Dermacentor variabilis*^8,9^. A synganglion transcriptome was also sequenced and annotated for *Ixodes scapularis*^6^. The above studies have identified several genes associated with the neural system and nerve impulse transmission, for example neuropeptides and proteins associated with neurotransmission. The synganglion transcriptome of the cattle tick, *Rhipicephalus microplus*, was investigated with a focus on predicting genes in the G protein-coupled receptor (GPCR) family^10^, representing potential targets for new acaricides^11^. But in each of the above studies, relatively few genes associated with neurotransmission were reported, probably due to relatively low sequencing depth. Further, in these data sets, it is impossible to assess the extent to which these gene collections were synganglion specific, as expression levels were only measured for synganglion samples, and were not compared to other tissues or the whole body. Recently, the *Ixodes scapularis* ISE6 cell line was shown to have neuronal characteristics, which provided a base for investigating neuronal interactions between ticks and pathogens^3^.

Ligand-gated cys-loop ion channels (cys-loop LGICs) are key receptors mediating neurotransmission both in vertebrates and invertebrates. Ligand-gated ion channels are major pharmacological targets of pesticides or acaricides such as macrocyclic lactones, phenylpyrazoles and neonicotinoids ^5^. Yet, their knowledge for ticks is partial in several aspects, including sequence reconstruction and annotation, phylogeny, or function, even though complete genomes of ticks have been sequenced for *Ixodes scapularis* ^4^ (32 sequences of cysLGICs were found in this study) or five other tick species^12^ (no specific annotation of cysLGICs reported in this study). The cys-loop LGICs form a superfamily of subunits that share a similar membrane topology. Indeed, they comprise one large amino terminal ligand-binding domain and four transmembrane domains, the second of these domains determining the ionic pore. The ionic pore consists of a multimeric assembly of five subunits, which can be identical or different. Subgroups within LGICs are determined by the activating neurotransmitter molecules - acetylcholine, *γ*-Aminobutyric acid (GABA), glycine, glutamate, or histamine - able to bind and trigger the opening of the channel gate. In Chelicerata, a first extensive reconstruction and in-depth study of the cys-loop LGICs was carried out for the two-spotted spider mite *Tetranychus urticae* (Acari: Acariformes), a phytophagous parasite, on the basis of its genome sequence^13^. On the basis of its transcriptomes, a similar reconstruction and phylogenetic study was carried out for the spider *Cupiennius salei*^14^. Similar analysis is lacking for ticks, despite the important implications of this knowledge for a better understanding of tick neurobiology and for providing potential targets for their control. The tick *Ixodes ricinus* is a widespread species in Europe, where it represents an increasing human health concern, being the main potential transmitter of the Lyme disease agent and of other microbes or parasites^15^. No transcriptome of the synganglion has been sequenced and no annotation of the cys-loop LGICs has been yet performed for this species, to our knowledge.

To study the interactions between ion channels and pesticides, i.e. between the nicotinic receptors of acetylcholine (nAChRs) and neonicotinoids, and between GABA receptors and other molecules, and to evaluate possible adverse effects on non-target species, systems of *ex-vivo* expression constitute an approach of choice. However the heterologous expression of some ion channels such as cholinergic receptors remains challenging. This has been succesful in some arthropods using the alpha subunits of the target species combined with vertebrate beta subunits (making hybrid receptors), whereas a first non-hybrid arthropod receptor using alpha and beta units from the target species, was obtained for the salmon louse *Lepeophtheirus salmonis*^16^. For arachnids, the first functional characterization of an alpha-type receptor was done for the brown dog tick *Rhipicephalus sanguineus*, but in vitro expression patterns were unreliable^17^. In the tick *I. ricinus*, the microtransplatation of synganglion membranes in oocytes of *Xenopus levis* allowed to express nicotinic acetylcholine receptors and to evaluate their sensitivity to insecticides^18^, but insofar no cloning of alpha-type receptor has been reported for ticks. A GABA receptor was first identified in *Drosophila melanogaster* (the *Rdl* gene^19^, confering resistance to the insecticide dieldrin), whereas *Rdl* homologs were cloned in the ticks *Dermacentor variabilis* ^20^ and *Rhipicephalus microplus* ^21^ respectively.

In the present study, our principal aims were i) to characterize genes that are specifically expressed (or up-regulated) in the synganglion of the tick *Ixodes ricinus* ii) to undertake a test of ion channel functionality, using the *Rdl* gene as a model iii) to carefully reconstruct the sequences of the *Ixodes ricinus* cys-loop LGICs, and to study their phylogeny including homologs in Arachnida and other arthropods.

## Material & methods

### Tick sampling

Ticks (*Ixodes ricinus*) were collected under two conditions, unfed or partially fed. Unfed adult female and male ticks were collected by the flagging method on vegetation in Carquefou (Loire-Atlantique, France). Partially fed adult females (half-fed) were collected in Chizé (Deux-Sèvres, France), from roe deers captured in the framework of a long-term monitoring programme of these mammals in the Chizé Forest, a biosphere Reserve.

### Ethics statement

Roe deers were captured and released alive, following a protocol that complies with Directive 2010/63/EU of the European Parliament and of the council of 22 September 2010 on the protection of animals used for scientific purposes. This protocol was approved by the Ethics Committee in Animal Experimentation of the University Claude Bernard Lyon 1 (CEEA-55; DR2014-09). The capture of roe deer was carried out with minimal stress and pain to animals, which were released alive at the same site and on the same day of capture.

### Sample preparation

Ticks were dissected under a stereomicroscope within 2-3 days of collection. Whole ticks were washed in RNase-ExitusPlus™ plus water. Synganglia were excised, rinsed in PBS buffer 1X, and dried by placing them gently on a clean sheet of paper. Synganglia were then placed in RNase-free microfuge tubes containing RNA extraction buffer (RA1 buffer, Nucleospin RNA XS kit, Macherey Nagel) and stored at -80°C until use. For RNA extraction, the tubes were gently thawed and tissues were mechanically disrupted with decontaminated pestles (3%H2O2) and eluted in 200 microliters of RA1. Samples were prepared for three conditions, unfed females, males, and partially fed females, each with three replicates (Table 1). Samples were made with pools of synganglia (n=25 for unfed females or males, n=10 for partially fed females). RNA was extracted using the Nucleospin RNA XS kit (Macherey Nagel, Düren, Germany), adapted for small samples, with a 5U DNAse treatment.

**Table 1:** List of libraries used for the comparison of expression levels in the synganglion and in the whole body. Accession: Accession numbers of reads (synganglion libraries, generated for the present study, were deposited as BioProject PRJEB40724, whereas whole body samples correspond to BioProject PRJNA395009, produced in a previous work of our group^28^). Feeding condition: ticks were either unfed (collected on the vegetation) or fed (semi-engorged state of repletion, ticks collected on roe deers for the synganglion libraries and on mice or rabbits for the whole body libraries). Stage/sex: indicates if ticks were female or male (for adults), or nymphs (the sex of nymphs could not be determined). Tissue: libraries were obtained from the dissected synganglion (this study) or from tick whole bodies (already published sequence data).

| Library name | Accession | Feeding Condition | Stage/sex | Tissue |
| --- | --- | --- | --- | --- |
| F01 | SAMEA7383684 | Unfed | females | Synganglion |
| F02 | SAMEA7383685 | Unfed | females | Synganglion |
| F03 | SAMEA7383686 | Unfed | females | Synganglion |
| M01_02 | SAMEA7383691 | Unfed | males | Synganglion |
| M03 | SAMEA7383687 | Unfed | males | Synganglion |
| FSG01 | SAMEA7383688 | Fed | females | Synganglion |
| FSG02 | SAMEA7383689 | Fed | females | Synganglion |
| FSG03 | SAMEA7383690 | Fed | females | Synganglion |
| N1 | SRR5839703 | Unfed | nymphs | Whole body |
| N2 | SRR5839701 | Unfed | nymphs | Whole body |
| M1 | SRR5839697 | Unfed | males | Whole body |
| M2 | SRR5839696 | Unfed | males | Whole body |
| F1 | SRR5839704 | Unfed | females | Whole body |
| F2 | SRR5839708 | Unfed | females | Whole body |
| F3 | SRR5839707 | Unfed | females | Whole body |
| NG1 | SRR5839700 | Fed | nymphs | Whole body |
| NG2 | SRR5839699 | Fed | nymphs | Whole body |
| NG3 | SRR5839698 | Fed | nymphs | Whole body |
| FG | SRR5839710 | Fed | females | Whole body |

### RNA dosage and quality

RNA purity was assessed with a NanoDrop (Thermo Fisher Scientific Inc., Watham, MA, USA). The commonly used threshold to consider samples as sufficiently pure is A280/A260 >1.80. Three out of nine samples were just below that threshold (F03, M01 and M02) but were kept for further analyses. Quantity was assessed with a Qubit (Invitrogen, Thermo Fisher Scientific Inc., Watham, MA, USA): quantities were low in the three samples mentioned above (they were not even dosable), but ranged between 68 and 304 ng for other samples. Finally, a further assessment of quantity and quality was conducted with the 2100BioAnalyzer instrument (Agilent Technologies Inc., Santa Clara, CA, USA): quality was determined by peak profiles, which was good for all samples, with a single, clear peak corresponding to rRNA, but low quantities for three samples, in particular M01 and M02. We decided to pool the latter two samples to make a single library, named M01_02.

### Library preparation and sequencing

RNA-Seq library preparations were carried out from 60ng total RNA, or the total volume when not quantifiable (for F03, M01_02). We used the NEBNext Ultra II directional RNA library prep kit (NEB, Ipswich, MA, USA), which allows mRNA strand orientation (sequence reads occur in the same orientation as anti-sense RNA). Briefly, poly(A)+ RNA was selected with oligo(dT) beads, chemically fragmented and converted into single-stranded cDNA using random hexamer priming. Then, the second strand was generated to create double-stranded cDNA. cDNA were then 3’-adenylated, and Illumina adapters were added. Ligation products were PCR-amplified. Ready-to-sequence Illumina libraries were then quantified by qPCR using the KAPA Library Quantification Kit for Illumina Libraries (KapaBiosystems, Wilmington, MA, USA), and libraries profiles evaluated with an Agilent 2100 Bioanalyzer (Agilent Technologies, Santa Clara, CA, USA). Each library was sequenced using 151 bp paired end reads chemistry on a Illumina HiSeq 4000 sequencer, producing a minimum of 40 million paired-reads for each library.

### Read cleaning and assembly

Short Illumina reads were bioinformatically post-processed as in ^22^ to filter out low quality data. First, low-quality nucleotides (Q < 20) were discarded from both read ends. Then remaining Illumina sequencing adapters and primer sequences were removed and only reads ≥ 30 nucleotides were retained. These filtering steps were done using in-house-designed software based on the FastX package (https://www.genoscope.cns.fr/fastxtend). Finally, read pairs mapping to the phage phiX genome were identified and discarded using SOAP aligner ^23^ (with default parameters) and the Enterobacteria phage PhiX174 reference sequence (GenBank: NC_001422.1). In addition, ribosomal RNA-like reads were detected using SortMeRNA ^24^ and filtered out. This set of cleaned reads were deposited to EBI under the BioProject accession PRJEB40724. The quality of cleaned reads was evaluated with fastQC, using MultiQC for visualization^25^, all parameters being satisfactory. Potential duplicates were removed with the the tool fastuniq (https://sourceforge.net/p/fastuniq/). The assembly was done with Trinity (version 2.3.2)^26^ with the options normalize-reads and RF (to account for strandedness). Calculations were done on the Genotoul Bioinformatics platform, using 100GB of memory. We made two separate assemblies, for respectively unfed ticks (females and males) and feeding ticks (females only). This choice was motivated by the expectation of different profiles of transcription in both conditions^27^ and of a reduced risk of producing chimeric contigs compared to a single assembly of all the reads.

### Construction of a high quality reference transcriptome

The two assemblies produced for the synganglion transcriptomes were combined with two previously published transcriptomes of the tick whole body - assembly accessions GFVZ^28^ and GIDG^29^ in the TSA division of Genbank - to produce a reference meta-transcriptome. The rationale for mixing different sources of transcribed sequences was to obtain the most complete transcriptome possible, in order to better evaluate the specificity of transcripts in the synganglion versus all tissues combined (the whole body). We did not use other transcriptomes published for specific organs of *I. ricinus*, because they have been sequenced a relatively low coverage and show comparatively low completeness^28^. To obtain the meta-transcriptome, we used the DRAP software (RunMeta)^30^, a dedicated tool able to fuse contigs of different assemblies in a unique representative contig set. This tool allows to reduce redundancy within and between sets, and to eliminate low quality contigs with low read support. Completeness of the different transcriptome assemblies was assessed with BUSCO (version 4.0.2)^31^, using the arachnida_odb10 set of n=2,934 conserved genes.

### Transcriptome annotation

Annotation of the reference transcriptome was done following the Trinotate pipeline (https://github.com/Trinotate/Trinotate.github.io)^32^. First, genes were predicted with TransDecoder.Predict (https://github.com/TransDecoder/TransDecoder/wiki), a prediction guided by PFAM-A searches with hmmscan (to recognize conserved domains) and a homology search with Diamond^33^ blastp (https://github.com/bbuchfink/diamond) between long peptides and the uniprot_sprot database. Predicted peptides were then searched with SignalP (to determine the presence of signal peptides), tmhmm (to determine if transmembrane domains are present) and rnammer (to identify rRNA contigs). We also performed homology searches against the UniRef90 database, with Diamond blastp (for predicted peptides) and Diamond blastx (for transcripts). Finally an annotation report was generated to combine all results, including retrieved Gene Ontology terms, KEGG, Eggnog and Pfam domains.

### Annotation of cys-loop LGIC genes in the reference transcriptome

A collection of reference cys-loop LGIC proteins comprising genes from *D. melanogaster* and the mite *T. urticae*^13^ (Acari, Acariformes) was established, to be used as tblastn queries on the reference meta-transcriptome of *I. ricinus*. A careful examination of the matches was then performed, to remove any remaining redundancy. Based on similarities within contigs and preliminary phylogenetic analyses, we identified events of chimerism among different gene sequences, which occured in the GABA and nAChR families respectively. For these genes, we extracted matching reads, which were re-assembled with CAP3^34^, using stringent parameters. This allowed to disentangle chimerisms and reconstructing complete sequences within gene families. Once we achieved the annotation of cys-loop LGICs and occasional correction of the inital contigs, these genes were re-integrated in the meta-transcriptome: to avoid redundancy, the edited and curated sequences were used and substituted to the corresponding initial contigs, and were annotated with the same Trinotate pipeline.

### Read counts and Differential Expression Analyses

Alignment of reads with the meta-transcriptome including the curated cys-loop LGIC sequences were performed with Bowtie2(v2.4.1)-RSEM(v1.3.1), using default parameters. Expression levels were measured using the “count per million” metric produced by RSEM. Levels of expression were measured for eight libraries corresponding to the synganglion (produced for this study), but also, for comparison, for part of the “whole body” libraries used to build the reference meta-transcriptome as described above. For example, libraries from the GIDG transcriptome were not used because they were not replicated by condition. Read counts from the BioProject PRJNA395009 (GFVZ TSA assembly) were used, for all libraries except four shown to be of lesser quality in a previous study: library names and their accession are given in Table 1. First, to determine which transcripts were more expressed in the synganglion, we made a differential expression analysis (DE) comparing synganglion and whole body libraries, with edgeR and limma^35,36^. Second, to compare the patterns of expression in the synganglion of unfed and partially fed ticks (hereafter “unfed” and “fed” conditions), we used the read counts from the synganglion libraries only. The same methods were used in both DE analyses. We used the function gene_to_trans_map of edgeR, which assigned read counts to the same gene for contigs defined as isoforms based on their Trinity name (i.e. contigs that differed only by the final _i suffix). This resulted in 63194 unique genes. The statistics of each gene (read count, log-fold -change, significance, and annotation) were visually explored with the Glimma package^37^. Criteria for defining an up- or down-regulated gene were an absolute log-fold change >2 and an adjusted P-value (adj-P-value) < 0.05). The topGO package was used for the analysis of GO enrichment in genes up- or down-regulated in the synganglion (the whole body being taken as the reference) and for the analysis of enrichment in genes up- or down-regulated in the synganglion-”fed” libraries (the “synganglion-unfed” condition being taken as the reference) - (Alexa A, Rahnenfuhrer J (2021). *topGO: Enrichment Analysis for Gene Ontology*. R package version 2.44.0).

### Functional expression of tick neuroreceptors

The coding sequence of *Rdl* was PCR-amplified using pecific primers designed based on the synganglion transcriptome assembly (Iri-RDL-F0 GGTCAAGGAGGTCGCTTGCC; Iri-RDL-R0 ACGACAACTTTAAAGGCGAATGC, FuIri-RDL-ptb-XhoF AGCGATGGCGTTCAGTTGCTG; FuIri-RDL-ptb-ApaR GACCGTGTGCACTATTCGTCG). The *Rdl* PCR-product was subcloned into the transcription vector pTB-207 ^38^ using the In-Fusion® HD Cloning Kit (Clontech™) and cRNAs were synthesized with the T7 mMessage mMachine kit (Ambion™). Expression of GABA-Rdl in *Xenopus laevis* oocytes and two-electrode voltage clamp electrophysiology were carried out as described previously ^39^. Briefly, 36 nL of 700 ng/μL *Rdl* cRNAs were micro-injected into defolliculated *Xenopus* oocytes (Ecocyte Bioscience) using a Drummond Nanoject II microinjector and incubated 3 days at 19°C to allow subunit expression. Electrophysiological recordings were performed using an Oocyte Clamp OC-725C amplifier (Warner Instruments) under voltage clamp at -60 mV and analyzed using the pCLAMP 10.4 package (Molecular Devices).

### Phylogenetic analyses of the cys-loop LGICs

Because nAChRs constitute a divergent group within cysLGICs, we analysed separately this group of sequences. Thus, for respectively nAchRs and all other cysLGICs, we made aligments with MUSCLE, cleaned these alignments with GBLOCKs and further edited manually the alignments (with slight corrections to preserve conserved blocks). Phylogenetic maximum likelihood trees were obtained with IQ-TREE ^40^ the best model of substitution being determined with Model Finder ^41^, and branch support being assessed with 1000 ultrafast bootstrap replication ^42^. Phylogenetic trees were formatted graphically with ITOL^43^.

## Results

### Transcriptome assembly and reference transcriptome

After sequencing a total of 69.7 Gb, the reads of libraries corresponding to synganglions of ticks in “fed” and “unfed” conditions were assembled separately. Both assemblies were combined with two more assemblies recently obtained for whole ticks transcriptomes, to obtain a reference meta-transcriptome. Detailed assembly statistics for the meta-transcriptome and for its four components are given in Table 2. Numbers of contigs were >600K for the synganglion assemblies. The percentage of predicted complete BUSCO genes were relatively even among these four assemblies, ranging between 90.9 and 92.4%. The meta-transcriptome contained 70,125 contigs, a sharp decrease compared to the synganglion transcriptomes, which is explained by the filtering steps of Drap and the selection of contigs with a minimum read support, here 1 fpkm (Fragments Per Kilobase Million). The BUSCO completeness of the reference transcriptome (96.6%) was higher than any of the component assemblies, and the number of fragmented genes was very low (only 8).

**Table 2:** Statistics on transcriptome assemblies for *I. ricinus*, including two previously published datasets (transcriptomes of tick whole bodies), two transcriptome assemblies obtained for the tick synganglion (this study), and for the reference meta-transcriptome we obtained combining these assemblies, with Drap. Assembly short name (TSA accession for whole body transcriptomes), Tissue, stages, Read type (length and strandedness), Total sequenced bases, Number of contigs, N50 metric of each assembly, Maximum contig size, Busco completeness metrics, using the Arachnida odb10 data set (S, single and complete, D, duplicated and complete, F, fragmented, M, missing, %C, percentage of complete genes).

| Assembly | Tissue | Stages/Conditions | reads type | bases (Gb) | #contigs | N50 | max size | Busco completeness (Arachnida, odb10) |  |  |  |  |
| --- | --- | --- | --- | --- | --- | --- | --- | --- | --- | --- | --- | --- |
|  |  |  |  |  |  |  |  | S | D | F | M | %C |
| GFVZ | Whole Body | males, nymphs (fed or unfed), adult females (fed or unfed) | oriented, 2x125 bp | 20.10 | 179316 | 875 | 20233 | 2213 | 473 | 54 | 194 | **91.5** |
| GIDG | Whole Body | eggs, larvae, nymphs (fed or unfed) and adult females (fed or unfed) | non oriented, 2x50 bp | 21.52 | 55046 | 1979 | 24178 | 1852 | 816 | 38 | 228 | **90.9** |
| SYN-NG | Synganglion | males and adult females, unfed | oriented, 2x150 bp | 37.08 | 695933 | 523 | 21882 | 1518 | 1195 | 56 | 165 | **92.4** |
| SYN-FSG | Synganglion | adult females, fed | oriented, 2x150 bp | 32.59 | 602593 | 466 | 33016 | 1484 | 1203 | 58 | 189 | **91.6** |
| meta-transcriptome 1fpkm | - | - | - | 111.29 | 70125 | 1915 | 24178 | 2314 | 519 | 8 | 93 | **96.6** |

### Clustering of libraries based on read counts

We observed a strong clustering of libraries that corresponded to the same combination of tissue and feeding status, suggesting that other factors (stage, sex) influenced comparatively less the levels of expression (MDS plot, Figure 1). Indeed, the first axis of variation (axis 1, weight of 34%) clearly separated synganglion and whole body samples, whereas the second factor (axis 2, weight of 17%) separated the unfed and the partially fed conditions, respectively. Of note, expression levels were much less influenced by the feeding status in synganglion libraries, compared to whole body libraries. We also observe that the “fed-synganglion” libraries were somewhat less differentiated from whole-body libraries than the “unfed-synganglion” libraries (see Discussion). Given our focus on specific patterns of expression of the synganglion, we decided to make a differential expression analysis in two steps: first, we compared all synganglion libraries on one side versus all whole body libraries on the other side. Second, for the synganglion libraries only, we made a comparison betwen the “fed” and “unfed” samples, to evaluate the effect of the feeding status on expression levels in this tissue.

**Figure 1:**
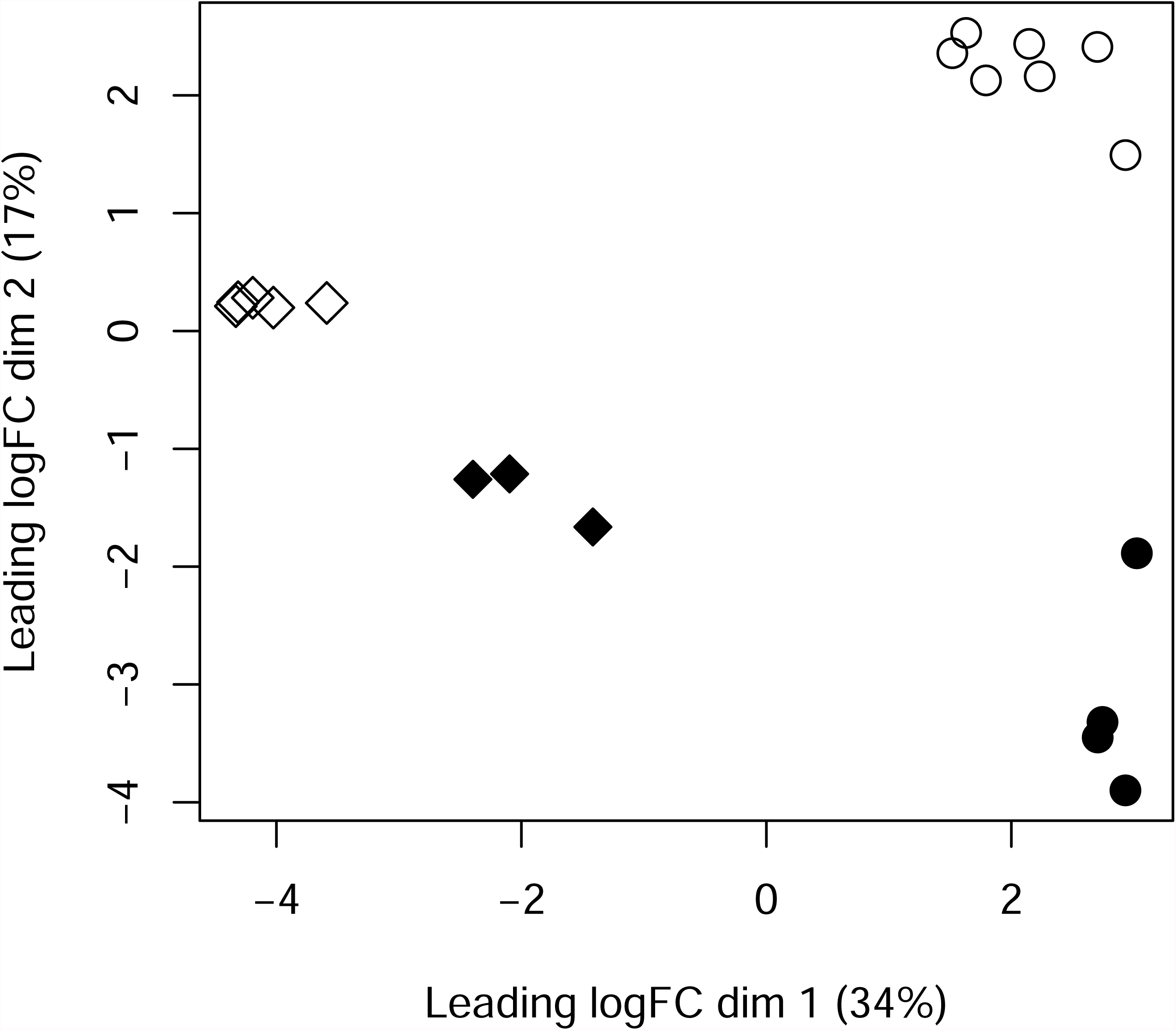
MDS plot, based on the read counts of libraries used for a differential expression analysis. Plot shown for dimensions 1 and 2, the two axes with the highest weight. Synganglion libraries were produced in this study, whole body libraries were from a previous study (as described in Table 1). Synganglion of unfed ticks (open diamonds), synganglion of fed ticks (filled diamonds), whole body of unfed ticks (open circles), whole body of fed ticks (filled circles).

### Differential expression analysis, comparison Synganglion vs Whole Body

A total of 8483 genes were up-regulated in the synganglion (Syn+ genes) while 5040 genes were down-regulated (Syn-genes), the remaining 22592 genes showing no significant difference between the synganglion and whole body samples. Therefore ∼37% of genes were found to be differently expressed among tissues. Many GO term associated with up-regulated genes for the Biogical Process (BP) and Molecular Function (MF) categories were consistent with neuronal functions (Table 3). This is the case of the terms GO:0098632 (cell-cell adhesion mediator activity), GO:1904315 (transmitter-gated ion channel activity involved in regulation of postsynaptic membrane potential, a GO term associated with cys-loop LGICs). As for the localization of expression (Cellular component, CC), many of the enriched terms were consistent with post-synaptic membranes or neurons. We also note the importance of terms associated with a large family of membrane-associated proteins, the G protein-coupled receptors (GPCRs). An MD plot also allowed to visualize genes with a significantly change of expression level in both tissues (Figure 2). Among genes with the highest over-expression in the synganglion, we note the cases of three genes of interest (Putative apoptosis-promoting rna-binding protein tia-1/tiar rrm superfamily, Putative beta-amyloid-like protein, Microtubule-associated protein tau). Most of the cys-loop LGICs were characterized by a strong over-expression in the synganglion (Figure 3): this included the GABA-1 (*Rdl*) receptor, and the eight nAChRs. We note that three of the nAChRs (beta1, alpha1 and alpha2) had very high and roughly equal levels of expression in the synganglion, suggesting a possible co-expression and that these three genes could form heteromeric receptors. By contrast, the levels of expression of some of the histamine receptors were not significantly different between the synganglion and whole body samples respectively, as was the case of Ig1, the homolog of CG7589, CG6927, and CG11340 (forming the clade Insect group 1, in the house-fly).

**Table 3:**
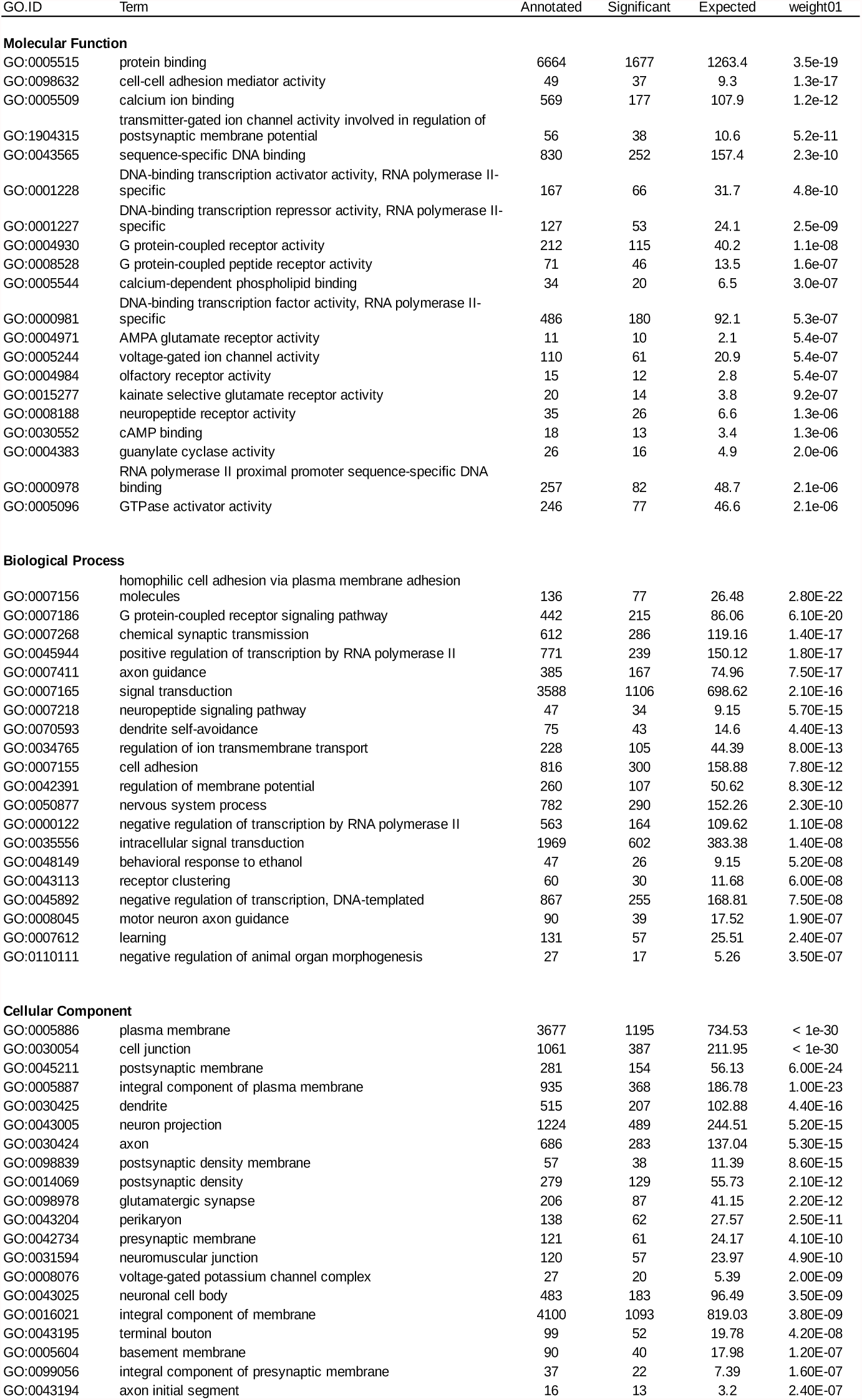
Top 20 enriched Gene Ontology terms among genes significantly up-regulated in the synganglion, compared to the whole body. Columns, GO IDs and terms for Molecular Function, Biological Process and Cellular localization; Annotated : number of transcripts with each GO; Significant: number of genes annotated with this term in the enriched category. Expected: expected number of genes. Weight01: level of significance of the enrichment.

**Figure 2:**
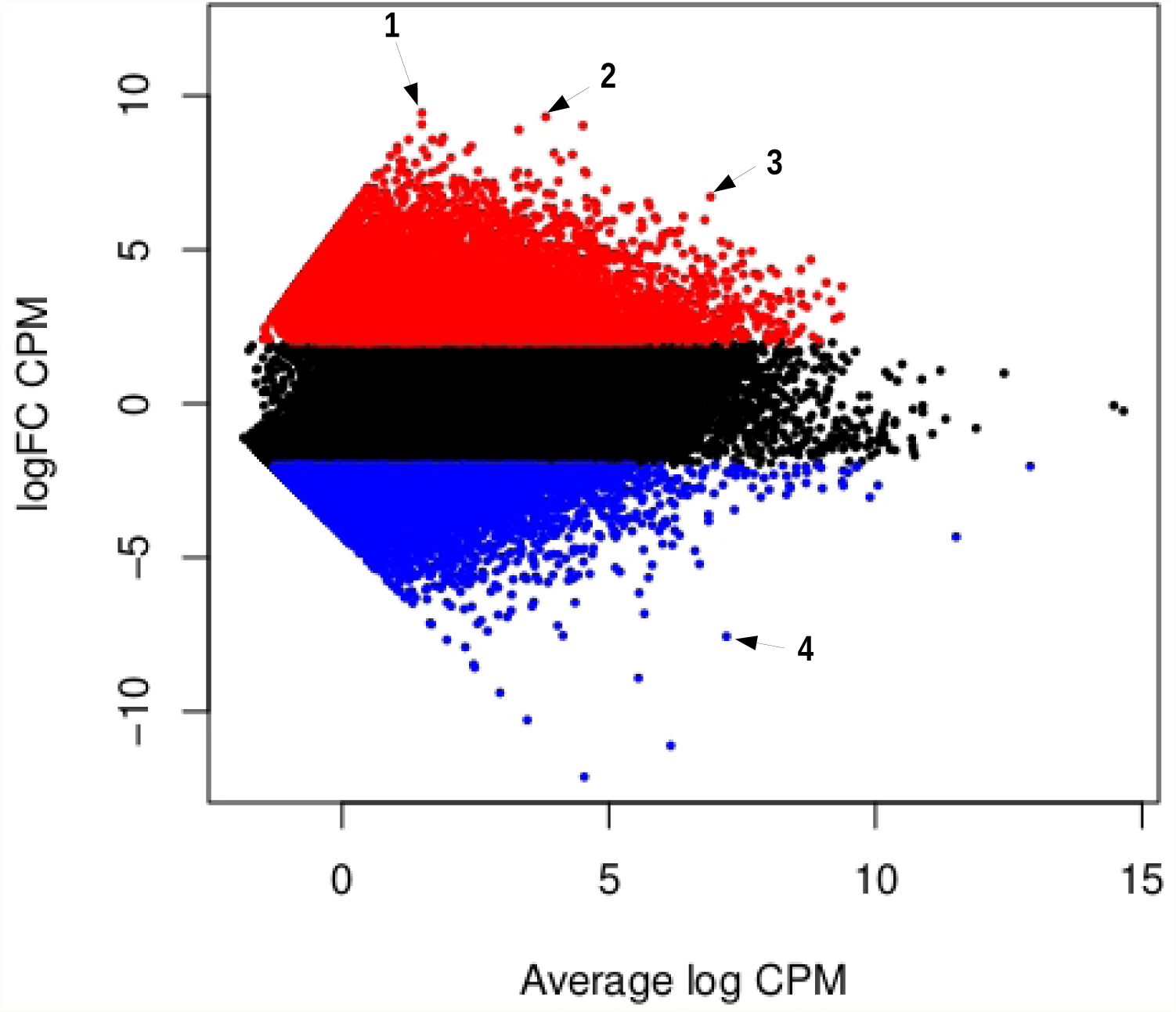
MD plot of all genes in the reference transcriptome, in a comparison of expression among synganglion and whole body libraries. In y-axis, log-fold change betwen the expression level in the synganglion and in the whole body. In x-axis: average log count per million of reads (averaged across all libraries, indicating the overall level of expression). Dots in red correspond to genes that are significantly up-regulated in the synganglion (logfold change >2 and P-adjusted value <0.05); dots in blue correspond to genes that are significantly down-regulated in the synganglion (logfold change <2 and P-adjusted value <0.05); dots in gray are genes with an expression level not signicantly different among tissues. Three outliers with among the highest values of logfold-change are distinguished (blastx description betwen parentheses): 1 (Putative apoptosis-promoting rna-binding protein tia-1/tiar rrm superfamily), 2 (Putative beta-amyloid-like protein), 3 (Microtubule-associated protein tau).

**Figure 3:**
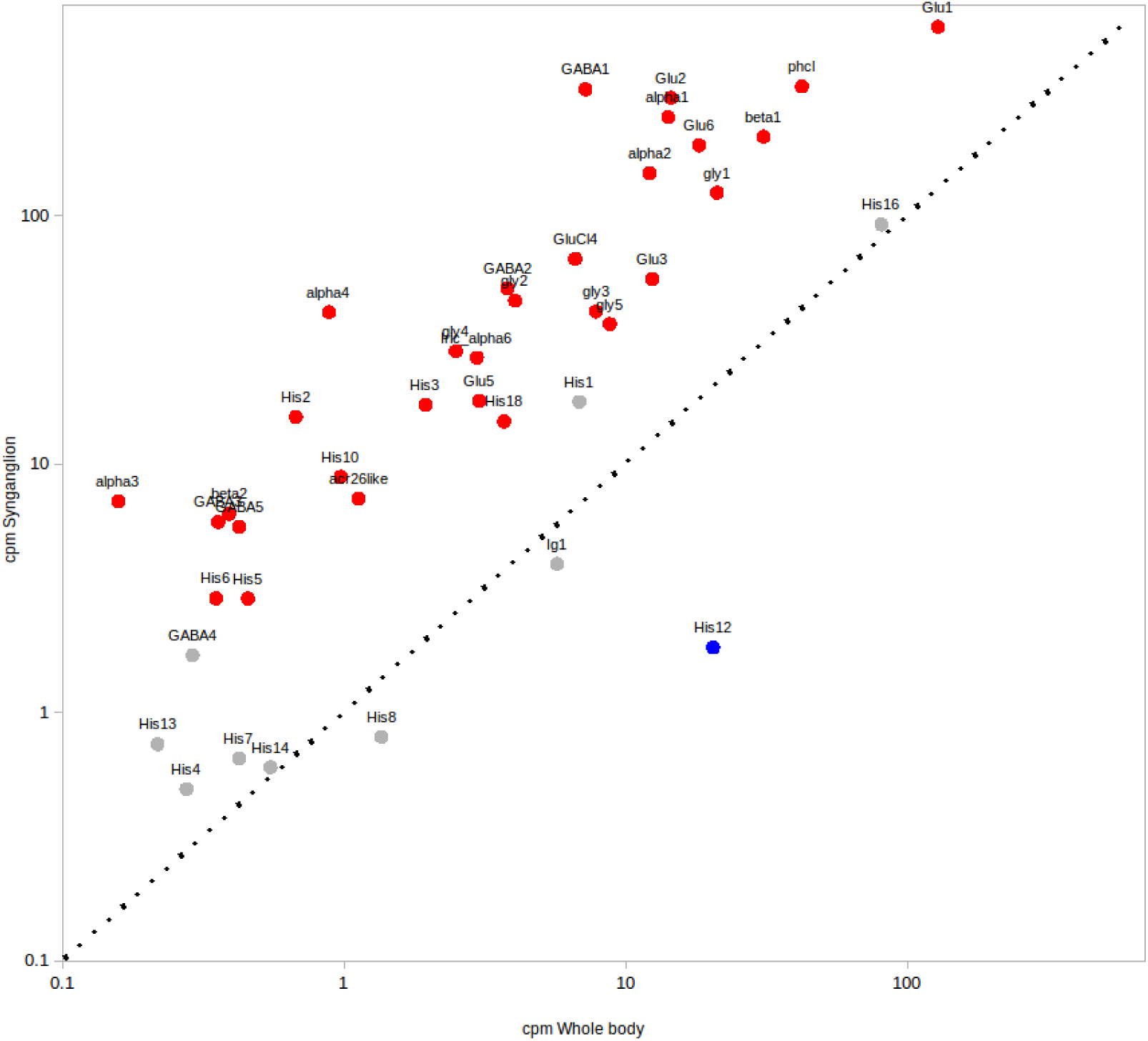
Comparison of expression counts, mesured in counts per million reads (cpm) with RSEM, for the cys-loop LCICs. Expression counts were averaged for all synganglion libraries (y-axis) and for all whole body libraries (x-axis). Colors of dots indicate if genes were found to be significantly up-regulated in the synganglion (red), down-regulated in the synganglion (blue) or with no sig. difference (grey). The dotted-line corresponds to y=x. For both axes, a logarithmic scale was used.

For the MF category, GO terms enriched in genes down-regulated in the synganglion included several linked terms (associated to the same genes): the first three categories (Iron ion biding, heme binding, aromatase activity) were indeed often found for genes belonging to the cytochrome P450 family (Supplementary Table 1).

### Comparison Fed versus Unfed for the synganglion libraries

A total of 2243 genes were up-regulated in the fed condition (Fed+ genes), whereas 2305 genes were down-regulated (Fed-genes) - for the remaining 35667 genes, the expressions levels did not differ significantly between the fed and unfed samples. This amounted to ∼11% of genes differently expressed among feeding conditions. The functional profiles of Fed+ genes showed notably an enrichment in genes associated with the metabolism of chitin and with the cuticle, whereas Fed-genes were enriched in a variety of GO terms, several of them being also enriched in the Syn+ category defined in the Synganglion vs Whole body comparison (e.g. terms associated with the synapse and with the plasma membrane). Among Syn+ genes, 1362 were Fed-whereas only 79 were Fed+.

### Functional expression of Rdl in Xenopus laevis oocytes

A full-length cDNA was obtained by RT-PCR, using the primers designed using the *Rdl* contig. The corresponding cRNA was thenmicroinjected into *Xenopus laevis* oocytes in order to monitor the potential functionnal expression of the RDL receptor. When challenged with 100 μM GABA, the micro-inected *Xenopus* oocytes responded with robust currents in the μA range (Figure 4A), providing a functional evidence of the expression of RDL. To further characterize this effect, the GABA concentration–response relationship was obtained by applying increasing concentrations of GABA (1 – 300 μM). The GABA concentration-response curve revealed an EC_50_ value of 19.95 μM (log EC_50_ = –4.7 ± 0.035, *n* = 18), with current amplitudes normalized to the maximal response obtained to 300 μM GABA (Figure 4B).

**Figure 4:**
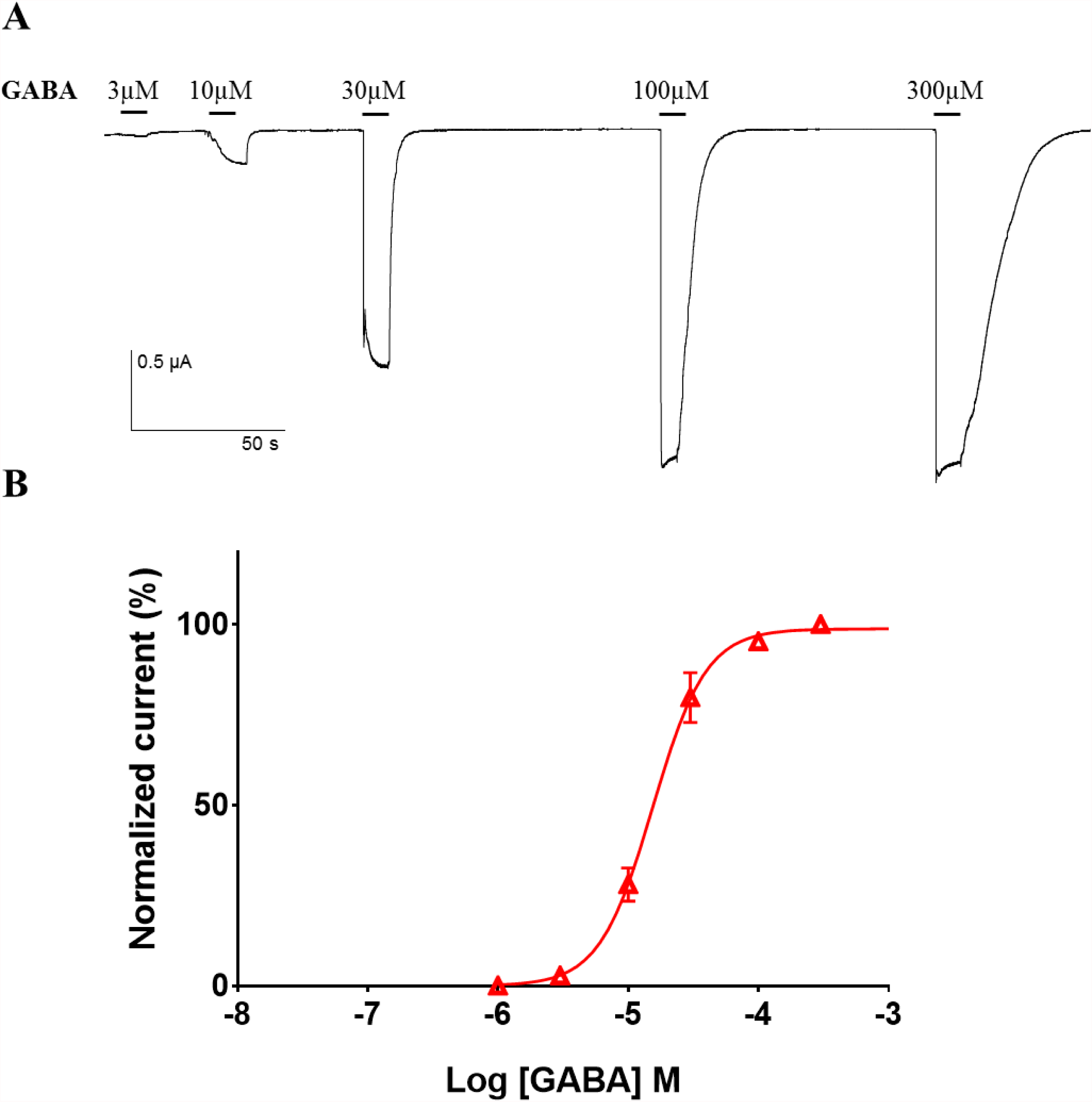
Concentration–response relationship of GABA on the *I. ricinus* RDL receptor expressed in *Xenopus* oocytes. A. Representative current traces of a single oocyte microinjected with *Iri-rdl* cRNA perfused with increasing concentrations of GABA for 10 seconds (short bars). The concentration of GABA (μM) is indicated above each trace. B. Concentration–response curve for GABA on the Iri-RDL channel. All current responses are normalized to 300 μM and shown as the mean ± SEM.

### Phylogeny of AChRs from ticks and other Arachnida

The ML phylogenetic study of tick nAChRs, including outgroups from other Arachnida and from an insect, *D. melanogaster*, allowed us to define eight clades with strong bootstrap support (Figure 5). Because genes of this family have been named and numbered independently in each species, and due to duplication or gene loss in different branches, a consistent naming system valid for all species is impossible, so we numbered clades based on orthogroups in Arachnida. Four clades named a1-4 were related to four genes in *D. melanogaster* (Dmela1, Dmela2, Dmela3 and Dmelb2), but there was no one-to-one orthology between the genes from Arachnida and from *D. melanogaster* respectively. For example the a1 and a2 clade were co-orthologous to Dmela1 whereas the a3 clade of Arachnida was orthologous to the pair of co-orthologs Dmela2 and Dmela3. The a6 and b1 clades were orthologous to *D. melanogaster* co-orthologs Dmela5-6-7 and Dmelb1, respectively. A second beta-type clade of Arachnida genes, named b2, was present both in ticks and other Arachnida but had no homologs in the genome of the house-fly nor in other insect genomes. This was also the case of a clade of alpha-type copies, that we named a5. Last, we identified a group of divergent nAChR genes (characterized by very long branches) comprising both the sequence Dmelb3 and sequences found in the spider *Parasteatoda tepidariorum*, but apparently without orthologs in ticks.

**Figure 5:**
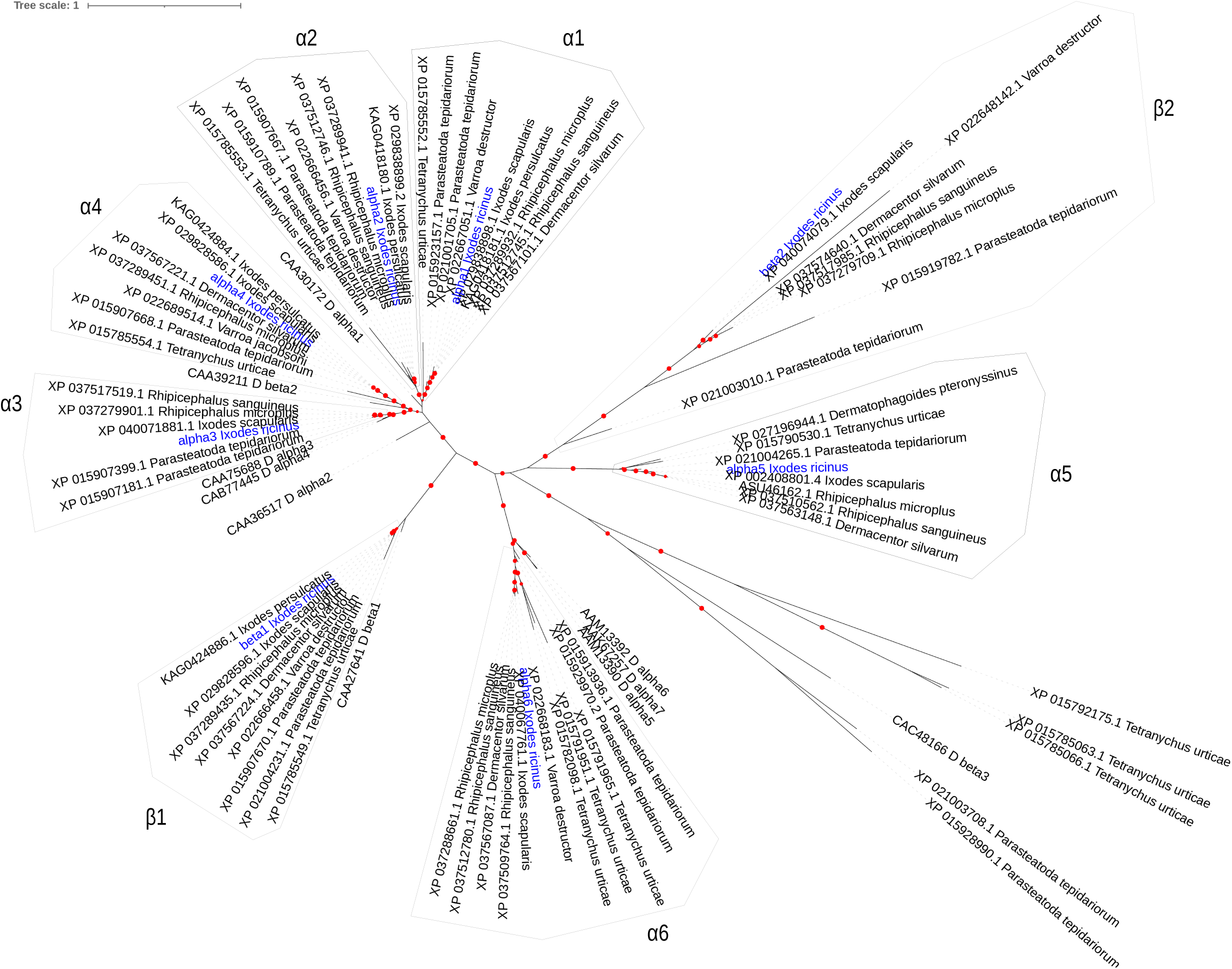
Maximum-likelihood phylogeny of nAchRs, including eight sequences identified in the synganglion transcriptome of the tick *I. ricinus* and their homologs in Arachnida (other tick species, other Parasitiformes, Acariformes) or in *Drosophila melanogaster*. Labels indicate the accession numbers of protein sequences, and species name (abbreviated as D for *D. melanogaster*). Labels for *I. ricin*us sequence are in blue. Filled circles on branches indicates bootstrap support (support increases with circle width, ranging from 80 to 100).

### Phylogeny of the non-AChR cysloop-LGICs

The numbers of different cysLGICs in *I. ricinus* and other representative arthropods are summarised in Table 4 : the total was 21 in the house-fly, while 54 genes were found in the house spider (*P. tepidariorum*) and 46 in *I. ricinus*. Based on our phylogenetic tree supported by high bootstrap values, we could define eight major clades of non-AChR cysloop-LGICs in Arachnida, comprising ticks and predatory mites (Parasitiformes, e.g. *Galendromus occidentalis*), plant-feeding mites (Acariformes, *e*.*g. Tetranychus urticae*) and spiders (Figure 6):

**Table 4:** Numbers of cys-loop LGICs in *D. melanogaster* and in three Chelicerata, the house spider *P. tepidariorum*, the mite spider *T. urticae* and the tick *I. ricinus*.

|  | *D. melanogaster* | *P. tepidariorum* | *T. urticae* | *I. ricinus* |
| --- | --- | --- | --- | --- |
| *Nicotinic Acetylcholine receptors (nAChRs)* |  |  |  |  |
| alpha1-2-3-4 group | 4 | 9 | 3 | 4 |
| beta1 group | 1 | 2 | 1 | 1 |
| alpha6 group | 3 | 4 | 3 | 1 |
| beta2 group | 0 | 2 | 0 | 1 |
| acr26-like group | 0 | 1 | 1 | 1 |
| Divergent nAChRs | 1 | 5 | 2 | 0 |
| *GABA-gated ion channels* |  |  |  |  |
| Rdl | 1 | 1 | 3 | 1 |
| Grd | 1 | 1 | 0 | 1 |
| Lcch3 | 1 | 2 | 0 | 1 |
| CG8916 | 1 | 1 | 0 | 1 |
| Other | 0 | 2 | 0 | 1 |
| *Other Cys-loop LGICs* |  |  |  |  |
| Histamine-gated | 2 | 10 | 4 | 18 |
| Group 12344 | 1 | 0 | 4 | 0 |
| Glutamate-gated | 1 | 5 | 6 | 6 |
| pHCl | 1 | 3 | 1 | 1 |
| Insect group1 | 3 | 2 | 0 | 1 |
| Glycine group | 0 | 4 | 1 | 5 |
| Total cys-loop LGICs | **21** | **54** | **29** | **44** |

**Figure 6:**
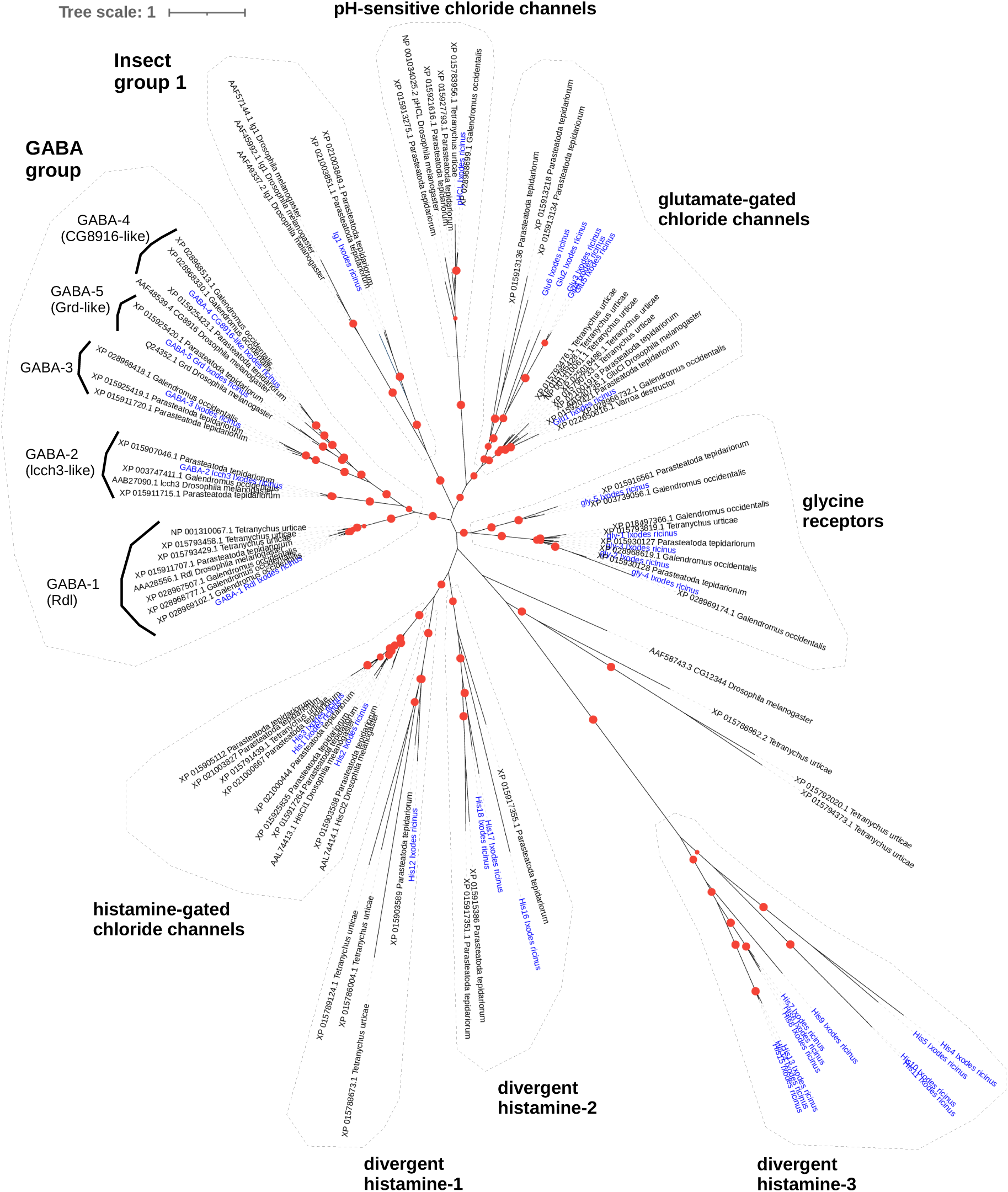
Maximum-likelihood phylogenetic tree of cys-loop LGICs subunit protein sequences, excluding nACHRs, in the tick *I. ricinus* and representative arthropods: *P. tepidariorum* (house spider), Acariformes and Parasitiformes, *D. melanogaster*. Labels indicate accession numbers of protein sequences and species name. Labels for *I. ricin*us sequence are in blue. Filled circles on branches indicates bootstrap support (support increases with circle width, ranging from 80 to 100).

i. GABA-group: five different sub-groups were found, including GABA-1, corresponding to the “Resistant to dieldrin” (Rdl) homologs, GABA-2, corresponding to “ligand-gated chloride ion channel homolog 3” (lcch3), GABA-3 which was represented in several Arachnida groups, but not in *D. melanogaster*, GABA-4, corresponding to homologs of the CG8916 gene in *D. melanogaster*, and GABA-5, representing “GABA and glycine-like receptor” (grd) homologs. Of note, Rdl in *I. ricinus* was characterized by three alternative isoforms: their sequences were identical except in a stretch of 154 amino acids, which did align well but showed several substitutions, suggesting the presence of alternative exons (alignment shown in Suppl. Figure 1).
ii. pH-sensitive chloride channels (pHCl): a single homolog was identified for *I. ricinus*, compared to three in *P. tepidariorum*.
iii. “insect group 1”: this clade comprised three copies in *D. melanogaster*, with one homolog in *I. ricinus* and two in *P. tepidariorum*.
iv. Glutamate gated chloride channels: within this group, we could distinguish one “conserved” subclade - one homolog in *I. ricinus* (Glu1) but multiple copies in *P. tepidariorum* (2) or *T. urticae* (5) - and one “divergent” subclade represented in *I. ricinus* (5 copies) and *P. tepidariorum* (3 copies).
v. Glycine receptors were represented by multiple copies in several groups of Arachnida, with 5 copies in *I. ricinus*.
vi. Histamine-gated chloride-channels (HisCl) : three copies in *I. ricinu*s (His1, His2 and His3) were found to be homologous of the genes HisCl1 and HisCl2 in *D. melanogaster*.
vii. other clades: we found three more clades represented only in Arachnida, or only in ticks, all characterized by long branches, i.e. by divergent sequences. This was the case of a clade with one copy in *I. ricinus* (His 12): this clade grouped robustly with the Histamine clade, so we named it “Divergent histamine 1”. A second clade, related to these genes, but with lower bootstrap support (<80) comprised genes in *I. ricinus* (His16-17-18), and also three genes in *P. tepidariorum* : we named it “Divergent histamine 2”. The third clade, characterized by particularly divergent sequences, comprised as many as 14 copies in *I. ricinus* (named His4 to His15) for which we found no homolog in other Arachnida. In our phylogeny, this group branched with the gene CG12344 from *D. melanogaster*, but with low bootstrap support. Based on the genomic organization in *I. scapularis* (Supplementary Figure 2) we found that eight of these copies (His4-His11) formed a cluster of tandem copies immediately adjacent to genes of the conserved HisCl clade (*I. scapularis* sequences homologous to His1, His 2 and His3). This very close physical proximity suggest that all of these copies probably emerged through tandem duplication, and that the digergent clade (His4 to His 11 and His 13 to His15) and His-gated subunits have a common ancestor. We then named it “Divergent histamine 3”.

## Discussion

### A meta-transcriptome of high completeness, enriched by synganglion transcriptome sequencing

Our work is based on the high throughput sequencing of a synganglion transcriptome, a first for the tick *Ixodes ricinus*, but also on the combination of these data with two previous assemblies of tick whole body transcriptomes. This allowed us to obtain a meta-transcriptome of very high completeness (>96% BUSCO completeness), an essential step for comparing expression levels in the synganglion and in the whole body. This approach eliminated a large part of the noise associated with *de novo* RNA-Seq assemblies (our two assembled synganglion transcriptomes each had >600K contigs), as we obtained a compact reference assembly with ∼70K contigs only. This reduction, due to the filtering of contigs with very low read support, could be associated with a loss of information, but high completeness metrics of the meta-transcriptome indicate that this risk was reduced. Overall, it appears that our newly acquired transcriptomic data for the synganglion has greatly improved the level of completeness of the meta-transcriptome, thanks to the inclusion of many transcripts that are highly specific to the synganglion.

### High expression specificity and neuronal signature of the synganglion

The DE study comparing synganglion and whole body samples showed a strong association between genes up-regulated in the synganglion and neuronal/neurotransmission functions. Still, a possible limitation of our approach is that the tissues we dissected and determined to be the synganglion could have been contaminated by adjacent tissues or cells, for example due to the particular organisation of this tissue which is crossed by the oesophagus and surrounded by the fat body. Although it is difficult to rule out this possibility entirely, the high specificity and distinct functional profile of the synganglion libraries indicate that contamination must have been limited. Enriched categories in the synganglion include neuropeptides, ion channels, etc., many of which were either completely absent or represented by partial sequences in transcriptomes obtained from the whole body or other tissues. A large majority of cysloop-LGICs were also upregulated in the synganglion, which is expected for these ion channels known to be expressed in neurons, which are particularly concentrated in the CNS. This overexpression concerns even divergent sequences of the cysLGICs (in the GluCls, or His-gated groups expanded clades), suggesting that these subunits are active mainly in the tick CNS. For the His-gated group, however, we note the exception of His12 (“divergent Histamine-1” clade) which was downregulated in the synganglion. A few of the cysLGIC genes were not differentially expressed (Synganglion vs Whole body comparison), including the homolog of ‘Insect group 1’ (a group comprising CG7589, CG6927, and CG11340 in *D. melanogaster*). This is consistent with a study that showed that IG1 is non-neuronal in this species^44^.

Although our study focused primarily on one class of ion channels, the cys-loop LGICs, our assemblies (the synganglion assemblies as the meta-transcriptome) provide rich material for gene discovery in many functional groups associated with neurotransmission or neuronal functions in general. For example, we found that several of the genes with the highest log-fold variation in expression in the synganglion have vertebrates homologues which have been extensively studied for their association with Alzheimer’s Disease (AD): this is the case for the APP-like peptide (APPL) and of tau protein^45^, as well as a homolog of TIA-1, a gene associated with the development of tau protein^46^. Interestingly, a putative tau homolog was also identified in the synganglion transcriptome of another tick species, *Dermacentor variabilis*, where it has been reported to be strongly up-regulated upon initiation of blood feeding (unfed versus partially-fed ticks)^8^.

In addition to this, in our study, a contig identified as Cathepsin B was among the most strongly downregulated genes in the synganglion. Cathepsin B has been shown to have complex interactions with AD and to be involved in the degradation or production of Abeta, as reviewed in ^47^. In *D. melanogaster*, APPL has been shown to be restricted to the nervous system, and the authors hypothesized that a neural-specific function encoded by the APP gene has been selectively maintained along evolution^48^. In the tick synganglion, the strong up-regulation (tau, APP, TIA1) or down-regulation (CathB) of genes known to be involved in AD pathology in humans illustrates the interest of arthropod models to study the expression profiles of these genes, their interactions and their evolution.

### Profiles of downregulated genes in the synganglion

For a complete understanding of tick neurobiology, it is also interesting to assess which genes are downregulated in the synganglion. Genes in this category showed enrichment in terms associated with iron binding, heme binding, and aromatase activity, in particular. In a previous study^28^ where we investigated expression in the whole body of the tick, comparing fed and unfed conditions, genes downregulated during feeding (i.e. significantly more expressed in the whole body of unfed ticks than of fed ticks) were particularly enriched in the same first three terms. We found that these terms were co-occurring in the same genes, most of which of the cytochrome P450 (cytP450) family. This suggests that one group of cytP450 tends to be particularly expressed in the whole body of unfed ticks, but is expressed at a much lower level both in the whole body of partially fed ticks and in the synganglion of ticks of all conditions.

### Differentiation between the “fed” and “unfed” state

Our design allowed to compare the expression levels in the synganglion of respectively unfed ticks (males or females) and partially fed female ticks. The transcriptomes of these two conditions were comparatively less differentiated than the synganglion and whole body samples (11% and 36% of DE genes in the two comparisons, respectively). Up-regulated genes upon feeding (fed+ genes) were enriched in chitin-associated proteins, a result that appears surprising given the absence of cuticle in this tissue. In a previous transcriptome study of the tick whole body^28^, cuticle associated proteins were also the most up-regulated genes upon feeding, suggesting that an increased expression of cuticle-related products upon feeding is a general pattern. To determine the intersection of changes upon feeding (in the synganglion) and of genes of expression among tissues (synganglion vs whole body), we counted the genes that were both differentially expressed in the fed vs unfed comparison and Syn+: with this filtering, up-regulated genes in the fed condition were down to 79 (from 2243) whereas down-regulated genes were down to 1362 (from 2305). This suggests that extremely few genes up-regulated in the synganglion were Fed+, and then that the ‘cuticle’ signature of Fed+ genes is not specific to the synganglion, but is a global pattern in the tick body. By contrast, a relatively large fraction of genes were at the same time Syn+ and downregulated in the synganglion upon feeding. Consistently, Fed-genes had functional profiles relatively similar to Syn+ genes (which can be seen comparing Table 4 and Suppl. Table 3). Overall, this indicates that the expression profile of the synganglion of unfed ticks has a more distinct profile (compared to the whole body) than the synganglion of feeding ticks. We observed this precise pattern for most of the cys-loop LGIC gene (most of them being Syn+), since their expression levels were often comparatively lower in the fed samples than in the unfed samples, as was the case of GABA-1 (Rdl), Glu-6, PhCl for example.We propose that the synganglion overall activity could be lowered when ticks are fixed on a host and start to feed, due to the down-regulation of several functions associated with mobility or sensorial activities related to host-search, for example. This can be compared to another study of the synganglion of unfed, part-fed and replete ticks^8^: in this study, expressions profiles of the latter two stages were relatively similar, whereas the unfed state had the most dissimilar profile, and the “replete” stage showed a decline in expression and specificity of expression, which was linked to a possible shift from active stages of the tick life-cycle (host search, feeding, mating) to late stages (oviposition, senescence and death).

### Completeness of cys-loop LGICs

Our synganglion transcriptomes allowed us, thanks to a careful manual curation, to obtain a remarkably large and complete collection of cys-loop LGICs compared to previous studies, even when based on complete genome sequences: for example, a compilation of cys-LGICs in the *I. scapularis* genome^4^ reported a total of 32 genes, less than the 44 found in our study. In the recent study of five tick complete genomes^12^, no specific compilation of cys-LGICs was reported, but our phylogenetic studies and the alignments we obtained indicate that for all these species, the cysLGIC record is less complete than in our study, with either some genes entirely missing or having only partial sequences, possibly due to annotation issues. This difference may be explained by difficulties associated with assembling and annotating the complex genomes of ticks, and illustrates the high value of transcriptome data for the tick CNS.

## Conclusions

our new transcriptome data for the tick synganglion, thanks to its quality and high coverage, allowed us to build a high quality reference transcriptome for *Ixodes ricinus*, strongly enriched in transcripts specific or at least strongly over expressed in neurones. This allowed us to reconstruct a large catalogue of the tick LGICs, showing that for the nAChRs and for the GABA group, this tick species has a repertoire comparable to other arthropods, while it shows marked expansions of other cysLGICs, especially the GluCls and the Histamine gated-like group. For GABA-Rdl, an internal duplication (triplication of an exon) allows the production of three alternative transcripts, either with or without a mutation confering resistance to dieldrin. Based on these synganglion transcriptomic ressources, we achieved the first functional reconstitution of a GABA-gated channel for *Ixodes ricinus*, made of the RDL subunit. Future studies will be needed to evaluate the pharmacological and biophysical properties of RDL and of other neuroreceptors, including sensitivity to insecticides.

## Acknowledgements

We thank Albert Agoulon (BIOEPAR, Nantes, France) for help in tick collection and Gilles Capron (OFB, Chizé, France), for giving us access to ticks collected on roe deers at the forest of Chizé. We also thank Ladislav Šimo (BIPAR, Maison-Alfort, France) for technical guidance in dissecting tick synganglia, Nicolas Lamassiaude (ISP, Tours, France) for valuable technical help on heterologous expression of ion channels, and Bioinfo Genotoul (Toulouse, France) for bioinformatic support.

## Availability of data

Read sequences of the synganglion transcriptomes have been published to EBI as BioProject PRJEB40721. The nucleotide sequences of cysLGICs have been deposited to GenBank as MZ027281-MZ027288 (n=8 nAChRs) and MZ099581-MZ099616 (other cysLGICs, n=36 transcripts, n=34 different genes). In addition, the gly1 and gly2 sequences correspond to accessions GIDG01020391.1 (gly1, coding sequence 1-1978) and GIDG01020392.1 (gly2, coding sequence 1-1993). Contig sequences of the meta-transcriptome, their annotation table, interactive plots of the Synganglion-vs-Whole body differential expression analysis, non-aligned and aligned protein sequences of the cysLGICs are available in a public repository (https://data.inrae.fr): Rispe, Claude, 2021, “Transcriptome of the the tick synganglion”, https://doi.org/10.15454/FGTIHR“.

## References

1. Smarandache-Wellmann, C. R. Arthropod neurons and nervous system. Curr. Biol. 26, R960– R965 (2016).

2. Lees, K. & Bowman, A. S. Tick neurobiology: recent advances and the post-genomic era. Invert. Neurosci. 7, 183–198 (2007).

3. Mateos-Hernández, L. et al. Enlisting the Ixodes scapularis Embryonic ISE6 Cell Line to Investigate the Neuronal Basis of Tick—Pathogen Interactions. Pathogens 10, 70 (2021).

4. Gulia-Nuss, M. et al. Genomic insights into the Ixodes scapularis tick vector of Lyme disease. Nat. Commun. 7, 10507 (2016).

5. Beugnet, F. & Franc, M. Insecticide and acaricide molecules and/or combinations to prevent pet infestation by ectoparasites. Trends Parasitol. 28, 267–279 (2012).

6. Egekwu, N., Sonenshine, D. E., Bissinger, B. W. & Roe, R. M. Transcriptome of the Female Synganglion of the Black-Legged Tick Ixodes scapularis (Acari: Ixodidae) with Comparison between Illumina and 454 Systems. PLoS ONE 9, (2014).

7. Lees, K., Woods, D. J. & Bowman, A. S. Transcriptome analysis of the synganglion from the brown dog tick, Rhipicephalus sanguineus. Insect Mol. Biol. 19, 273–282 (2010).

8. Bissinger, B. W. et al. Synganglion transcriptome and developmental global gene expression in adult females of the American dog tick, Dermacentor variabilis (Acari: Ixodidae). Insect Mol. Biol. 20, 465–491 (2011).

9. Donohue, K. V. et al. Neuropeptide signaling sequences identified by pyrosequencing of the American dog tick synganglion transcriptome during blood feeding and reproduction. Insect Biochem. Mol. Biol. 40, 79–90 (2010).

10. Guerrero, F. D. et al. Prediction of G protein-coupled receptor encoding sequences from the synganglion transcriptome of the cattle tick, Rhipicephalus microplus. Ticks Tick-Borne Dis. 7, 670–677 (2016).

11. Hill, C. A., Sharan, S. & Watts, V. J. Genomics, GPCRs and new targets for the control of insect pests and vectors. Curr. Opin. Insect Sci. 30, 99–106 (2018).

12. Jia, N. et al. Large-Scale Comparative Analyses of Tick Genomes Elucidate Their Genetic Diversity and Vector Capacities. Cell 182, 1328-1340.e13 (2020).

13. Dermauw, W. et al. The cys-loop ligand-gated ion channel gene family of Tetranychus urticae: Implications for acaricide toxicology and a novel mutation associated with abamectin resistance. Insect Biochem. Mol. Biol. 42, 455–465 (2012).

14. Torkkeli, P. H., Liu, H. & French, A. S. Transcriptome Analysis of the Central and Peripheral Nervous Systems of the Spider Cupiennius salei Reveals Multiple Putative Cys-Loop Ligand Gated Ion Channel Subunits and an Acetylcholine Binding Protein. PloS One 10, e0138068 (2015).

15. Medlock, J. M. et al. Driving forces for changes in geographical distribution of Ixodes ricinus ticks in Europe. Parasit. Vectors 6, 1 (2013).

16. Rufener, L., Kaur, K., Sarr, A., Aaen, S. M. & Horsberg, T. E. Nicotinic acetylcholine receptors: Ex-vivo expression of functional, non-hybrid, heteropentameric receptors from a marine arthropod, Lepeophtheirus salmonis. PLoS Pathog. 16, e1008715 (2020).

17. Lees, K. et al. Functional characterisation of a nicotinic acetylcholine receptor α subunit from the brown dog tick, Rhipicephalus sanguineus. Int. J. Parasitol. 44, 75–81 (2014).

18. Le Mauff, A. et al. Nicotinic acetylcholine receptors in the synganglion of the tick Ixodes ricinus: Functional characterization using membrane microtransplantation. Int. J. Parasitol. Drugs Drug Resist. 14, 144–151 (2020).

19. Ffrench-Constant, R. H., Mortlock, D. P., Shaffer, C. D., MacIntyre, R. J. & Roush, R. T. Molecular cloning and transformation of cyclodiene resistance in Drosophila: An invertebrate γ-aminobutyric acid subtype A receptor locus. Proc. Natl. Acad. Sci. U. S. A. 88, 7209–7213 (1991).

20. Zheng, Y., Priest, B., Cully, D. F. & Ludmerer, S. W. RdlDv, a novel GABA-gated chloride channel gene from the American dog tick Dermacentor variabilis. Insect Biochem. Mol. Biol. 33, 595–599 (2003).

21. Rufener, L., Danelli, V., Bertrand, D. & Sager, H. The novel isoxazoline ectoparasiticide lotilaner (Credelio™): a non-competitive antagonist specific to invertebrates γ-aminobutyric acid-gated chloride channels (GABACls). Parasit. Vectors 10, 530 (2017).

22. Alberti, A. et al. Viral to metazoan marine plankton nucleotide sequences from the Tara Oceans expedition. Sci. Data 4, 170093 (2017).

23. Li, R., Li, Y., Kristiansen, K. & Wang, J. SOAP: short oligonucleotide alignment program. Bioinformatics 24, 713–714 (2008).

24. Kopylova, E., Noé, L. & Touzet, H. SortMeRNA: fast and accurate filtering of ribosomal RNAs in metatranscriptomic data. Bioinforma. Oxf. Engl. 28, 3211–3217 (2012).

25. Ewels, P., Magnusson, M., Lundin, S. & Käller, M. MultiQC: summarize analysis results for multiple tools and samples in a single report. Bioinformatics 32, 3047–3048 (2016).

26. Haas, B. J. et al. De novo transcript sequence reconstruction from RNA-Seq: reference generation and analysis with Trinity. Nat. Protoc. 8, (2013).

27. Koči, J., Šimo, L. & Park, Y. Validation of Internal Reference Genes for Real-Time Quantitative Polymerase Chain Reaction Studies in the Tick, Ixodes scapularis (Acari: Ixodidae). J. Med. Entomol. 50, 79–84 (2013).

28. Charrier, N. P. et al. Whole body transcriptomes and new insights into the biology of the tick Ixodes ricinus. Parasit. Vectors 11, 364 (2018).

29. Vechtova, P. et al. Catalogue of stage-specific transcripts in Ixodes ricinus and their potential functions during the tick life-cycle. Parasit. Vectors 13, 311 (2020).

30. Cabau, C. et al. Compacting and correcting Trinity and Oases RNA-Seq de novo assemblies. PeerJ 5, e2988 (2017).

31. Seppey, M., Manni, M. & Zdobnov, E. M. BUSCO: Assessing Genome Assembly and Annotation Completeness. in Gene Prediction: Methods and Protocols (ed. Kollmar, M.) 227– 245 (Springer, 2019). doi:10.1007/978-1-4939-9173-0_14.

32. Bryant, D. M. et al. A Tissue-Mapped Axolotl De Novo Transcriptome Enables Identification of Limb Regeneration Factors. Cell Rep. 18, 762–776 (2017).

33. Buchfink, B., Xie, C. & Huson, D. H. Fast and sensitive protein alignment using DIAMOND. Nat. Methods 12, 59–60 (2015).

34. Huang, X. & Madan, A. CAP3: A DNA Sequence Assembly Program. Genome Res. 9, 868–877 (1999).

35. Robinson, M. D., McCarthy, D. J. & Smyth, G. K. edgeR: a Bioconductor package for differential expression analysis of digital gene expression data. Bioinformatics 26, 139–140 (2010).

36. Ritchie, M. E. et al. limma powers differential expression analyses for RNA-sequencing and microarray studies. Nucleic Acids Res. 43, e47–e47 (2015).

37. Law, C. W. et al. RNA-seq analysis is easy as 1-2-3 with limma, Glimma and edgeR. https://f1000research.com/articles/5-1408 (2018).

38. Boulin, T. et al. Eight genes are required for functional reconstitution of the Caenorhabditis elegans levamisole-sensitive acetylcholine receptor. Proc. Natl. Acad. Sci. 105, 18590–18595 (2008).

39. Lamassiaude, N. et al. The molecular targets of ivermectin and lotilaner in the human louse Pediculus humanus humanus: New prospects for the treatment of pediculosis. PLOS Pathog. 17, e1008863 (2021).

40. Nguyen, L.-T., Schmidt, H. A., von Haeseler, A. & Minh, B. Q. IQ-TREE: A Fast and Effective Stochastic Algorithm for Estimating Maximum-Likelihood Phylogenies. Mol. Biol. Evol. 32, 268–274 (2015).

41. Kalyaanamoorthy, S., Minh, B. Q., Wong, T. K. F., von Haeseler, A. & Jermiin, L. S. ModelFinder: fast model selection for accurate phylogenetic estimates. Nat. Methods 14, 587– 589 (2017).

42. Hoang, D. T., Chernomor, O., von Haeseler, A., Minh, B. Q. & Vinh, L. S. UFBoot2: Improving the Ultrafast Bootstrap Approximation. Mol. Biol. Evol. 35, 518–522 (2018).

43. Letunic, I. & Bork, P. Interactive tree of life (iTOL) v3: an online tool for the display and annotation of phylogenetic and other trees. Nucleic Acids Res. 44, W242–W245 (2016).

44. Remnant, E. J. et al. Evolution, Expression, and Function of Nonneuronal Ligand-Gated Chloride Channels in Drosophila melanogaster. G3 GenesGenomesGenetics 6, 2003–2012 (2016).

45. Bloom, G. S. Amyloid-β and tau: the trigger and bullet in Alzheimer disease pathogenesis. JAMA Neurol. 71, 505–508 (2014).

46. Apicco, D. J. et al. Reducing the RNA binding protein TIA1 protects against tau-mediated neurodegeneration in vivo. Nat. Neurosci. 21, 72–80 (2018).

47. Bernstein, H.-G. & Keilhoff, G. Putative roles of cathepsin B in Alzheimer’s disease pathology: the good, the bad, and the ugly in one? Neural Regen. Res. 13, 2100–2101 (2018).

48. Martin-Morris, L. E. & White, K. The Drosophila transcript encoded by the /J-amyloid protein precursor-like gene is restricted to the nervous system. 11.

