## Supplemental Tables and Figures for "Transcriptome of the synganglion and characterization of the cys-loop ligand-gated ion channel gene family in the tick *Ixodes ricinus*"

Supplementary Table 1 : Top 20 enriched Gene Ontology terms among genes significantly down-regulated in the synganglion, compared to the whole body. Columns, GO IDs and terms for Molecular Function, Biological Process and Cellular localization; Annotated : number of transcripts associated with each GO; Significant: number of genes annotated with this term in the enriched category. Expected: expected number of genes. Weight01: level of significance of the enrichment.

| GO.ID | Term | Annotated | Significant | Expected | weight01 |
|---|---|---|---|---|---|
| **Molecular Function** | | | | | |
| GO:0005506 | iron ion binding | 532 | 93 | 41.38 | 1.5e-13 |
| GO:0020037 | heme binding | 468 | 84 | 36.41 | 3.2e-13 |
| GO:0070330 | aromatase activity | 205 | 43 | 15.95 | 1.7e-09 |
| GO:0050649 | testosterone 6-beta-hydroxylase activity | 96 | 22 | 7.47 | 3.4e-06 |
| GO:0004222 | metalloendopeptidase activity | 168 | 31 | 13.07 | 5.3e-06 |
| GO:0015129 | lactate transmembrane transporter activity | 21 | 9 | 1.63 | 1.3e-05 |
| GO:0008401 | retinoic acid 4-hydroxylase activity | 63 | 16 | 4.9 | 1.8e-05 |
| GO:0016207 | 4-coumarate-CoA ligase activity | 22 | 9 | 1.71 | 2.0e-05 |
| GO:0101020 | estrogen 16-alpha-hydroxylase activity | 78 | 18 | 6.07 | 2.3e-05 |
| GO:0019825 | oxygen binding | 34 | 11 | 2.64 | 3.3e-05 |
| GO:0102756 | very-long-chain 3-ketoacyl-CoA synthase activity | 62 | 15 | 4.82 | 6.1e-05 |
| GO:0102336 | 3-oxo-arachidoyl-CoA synthase activity | 62 | 15 | 4.82 | 6.1e-05 |
| GO:0102337 | 3-oxo-cerotoyl-CoA synthase activity | 62 | 15 | 4.82 | 6.1e-05 |
| GO:0102338 | 3-oxo-lignoceronyl-CoA synthase activity | 62 | 15 | 4.82 | 6.1e-05 |
| GO:0004784 | superoxide dismutase activity | 23 | 8 | 1.79 | 0.00022 |
| GO:0050833 | pyruvate transmembrane transporter activity | 18 | 7 | 1.4 | 0.00025 |
| GO:0005496 | steroid binding | 106 | 17 | 8.25 | 0.00042 |
| GO:0004497 | monooxygenase activity | 438 | 78 | 34.07 | 0.00053 |
| GO:0005518 | collagen binding | 33 | 9 | 2.57 | 0.00071 |
| GO:0004252 | serine-type endopeptidase activity | 209 | 30 | 16.26 | 0.00081 |
| **Biological Process** | | | | | |
| GO:0006508 | proteolysis | 1655 | 149 | 129.54 | 1.6e-07 |
| GO:0055114 | oxidation-reduction process | 1361 | 166 | 106.52 | 2.3e-07 |
| GO:0055085 | transmembrane transport | 1325 | 125 | 103.71 | 3.6e-07 |
| GO:0051923 | sulfation | 89 | 22 | 6.97 | 9.5e-07 |
| GO:0008202 | steroid metabolic process | 541 | 72 | 42.34 | 7.4e-06 |
| GO:0009058 | biosynthetic process | 5826 | 451 | 456 | 8.9e-06 |
| GO:0070989 | oxidative demethylation | 110 | 23 | 8.61 | 1.1e-05 |
| GO:0009698 | phenylpropanoid metabolic process | 31 | 10 | 2.43 | 2.1e-05 |
| GO:0042381 | hemolymph coagulation | 29 | 10 | 2.27 | 4.2e-05 |
| GO:1901475 | pyruvate transmembrane transport | 13 | 6 | 1.02 | 5.3e-05 |
| GO:0035873 | lactate transmembrane transport | 7 | 5 | 0.55 | 5.4e-05 |
| GO:0046697 | decidualization | 12 | 6 | 0.94 | 0.00014 |
| GO:0019953 | sexual reproduction | 999 | 63 | 78.19 | 0.00026 |
| GO:0050790 | regulation of catalytic activity | 1263 | 72 | 98.85 | 0.00030 |
| GO:0051603 | proteolysis involved in cellular protein catabolic process | 797 | 50 | 62.38 | 0.00036 |
| GO:0042572 | retinol metabolic process | 80 | 16 | 6.26 | 0.00041 |
| GO:0006526 | arginine biosynthetic process | 6 | 4 | 0.47 | 0.00049 |
| GO:0061014 | positive regulation of mRNA catabolic process | 62 | 8 | 4.85 | 0.00049 |
| GO:0008209 | androgen metabolic process | 58 | 13 | 4.54 | 0.00055 |
| GO:0042359 | vitamin D metabolic process | 21 | 7 | 1.64 | 0.00077 |
| **Cellular Component** | | | | | |
| GO:0005576 | extracellular region | 2132 | 195 | 166.11 | 2.8e-06 |
| GO:0005764 | lysosome | 658 | 70 | 51.27 | 3.4e-05 |
| GO:0005840 | ribosome | 384 | 42 | 29.92 | 9.8e-05 |
| GO:0031932 | TORC2 complex | 14 | 5 | 1.09 | 0.0031 |
| GO:0005789 | endoplasmic reticulum membrane | 1406 | 140 | 109.55 | 0.0058 |
| GO:0005777 | peroxisome | 294 | 37 | 22.91 | 0.0085 |
| GO:0031526 | brush border membrane | 82 | 13 | 6.39 | 0.0105 |
| GO:0005739 | mitochondrion | 2170 | 193 | 169.07 | 0.0139 |
| GO:1905369 | endopeptidase complex | 115 | 11 | 8.96 | 0.0172 |
| GO:0042788 | polysomal ribosome | 22 | 5 | 1.71 | 0.0246 |
| GO:0030133 | transport vesicle | 349 | 14 | 27.19 | 0.0328 |
| GO:0046581 | intercellular canaliculus | 24 | 5 | 1.87 | 0.0349 |
| GO:0005779 | integral component of peroxisomal membrane | 41 | 7 | 3.19 | 0.0373 |
| GO:0009897 | external side of plasma membrane | 95 | 13 | 7.4 | 0.0398 |
| GO:0016324 | apical plasma membrane | 310 | 33 | 24.15 | 0.0484 |
| GO:0071797 | LUBAC complex | 11 | 3 | 0.86 | 0.0485 |
| GO:0035631 | CD40 receptor complex | 11 | 3 | 0.86 | 0.0485 |

Supplementary Table 2 : Top 20 enriched Gene Ontology terms among genes significantly up-regulated in the "fed" conditions, compared to the "unfed" condition, for synganglion libraries.

| GO.ID | Term | Annotated | Significant | Expected | weight01 |
|---|---|---|---|---|---|
| **Molecular Function** | | | | | |
| GO:0008061 | chitin binding | 159 | 33 | 4.99 | 3.6E-18 |
| GO:0042302 | structural constituent of cuticle | 97 | 24 | 3.04 | 2.1E-15 |
| GO:0051087 | chaperone binding | 108 | 16 | 3.39 | 0.00000025 |
| GO:0008453 | alanine-glyoxylate transaminase activity | 5 | 4 | 0.16 | 0.0000047 |
| GO:0051082 | unfolded protein binding | 156 | 17 | 4.89 | 0.0000084 |
| GO:0003998 | acylphosphatase activity | 6 | 4 | 0.19 | 0.000014 |
| GO:0004867 | serine-type endopeptidase inhibitor activity | 165 | 17 | 5.18 | 0.000018 |
| GO:0004252 | serine-type endopeptidase activity | 209 | 19 | 6.56 | 0.000034 |
| GO:0003796 | lysozyme activity | 8 | 4 | 0.25 | 0.000061 |
| GO:0032036 | myosin heavy chain binding | 18 | 5 | 0.56 | 0.00018 |
| GO:0008289 | lipid binding | 746 | 33 | 23.4 | 0.00023 |
| GO:0005125 | cytokine activity | 44 | 7 | 1.38 | 0.0004 |
| GO:0005200 | structural constituent of cytoskeleton | 107 | 11 | 3.36 | 0.00054 |
| GO:0008083 | growth factor activity | 60 | 8 | 1.88 | 0.00054 |
| GO:0098505 | G-rich strand telomeric DNA binding | 6 | 3 | 0.19 | 0.00057 |
| GO:0005212 | structural constituent of eye lens | 14 | 4 | 0.44 | 0.00075 |
| GO:0001640 | adenylate cyclase inhibiting G protein-coupled glutamate receptor | 9 | 3 | 0.28 | 0.00098 |
| GO:0016723 | oxidoreductase activity, oxidizing metal ions, NAD or NADP as ac | 8 | 3 | 0.25 | 0.00153 |
| GO:0004601 | peroxidase activity | 68 | 8 | 2.13 | 0.00162 |
| GO:0009922 | fatty acid elongase activity | 29 | 5 | 0.91 | 0.0019 |
| **Biological Process** | | | | | |
| GO:0006030 | chitin metabolic process | 95 | 21 | 2.89 | 7.5E-13 |
| GO:0042157 | lipoprotein metabolic process | 168 | 15 | 5.12 | 0.0000003 |
| GO:0060266 | negative regulation of respiratory burst involved in inflammatory | 21 | 7 | 0.64 | 0.0000019 |
| GO:0009436 | glyoxylate catabolic process | 5 | 4 | 0.15 | 0.0000042 |
| GO:0006937 | regulation of muscle contraction | 90 | 10 | 2.74 | 0.0000047 |
| GO:0002282 | microglial cell activation involved in immune response | 16 | 6 | 0.49 | 0.0000048 |
| GO:1905673 | positive regulation of lysosome organization | 16 | 6 | 0.49 | 0.0000048 |
| GO:0106016 | positive regulation of inflammatory response to wounding | 16 | 6 | 0.49 | 0.0000048 |
| GO:1902564 | negative regulation of neutrophil activation | 17 | 6 | 0.52 | 0.0000072 |
| GO:1905247 | positive regulation of aspartic-type peptidase activity | 17 | 6 | 0.52 | 0.0000072 |
| GO:1903979 | negative regulation of microglial cell activation | 17 | 6 | 0.52 | 0.0000072 |
| GO:1903334 | positive regulation of protein folding | 17 | 6 | 0.52 | 0.0000072 |
| GO:0009405 | pathogenesis | 17 | 6 | 0.52 | 0.0000072 |
| GO:0042026 | protein refolding | 36 | 8 | 1.1 | 0.00001 |
| GO:0002265 | astrocyte activation involved in immune response | 20 | 6 | 0.61 | 0.000021 |
| GO:1900426 | positive regulation of defense response to bacterium | 30 | 7 | 0.91 | 0.000026 |
| GO:0006636 | unsaturated fatty acid biosynthetic process | 62 | 9 | 1.89 | 0.000028 |
| GO:0042853 | L-alanine catabolic process | 7 | 4 | 0.21 | 0.000028 |
| GO:0007042 | lysosomal lumen acidification | 23 | 6 | 0.7 | 0.00005 |
| GO:0005975 | carbohydrate metabolic process | 703 | 32 | 21.42 | 0.000089 |
| **Cellular Component** | | | | | |
| GO:0005576 | extracellular region | 2132 | 157 | 67.16 | 4.3E-25 |
| GO:0005615 | extracellular space | 1146 | 60 | 36.1 | 0.0000084 |
| GO:0035578 | azurophil granule lumen | 46 | 8 | 1.45 | 0.000084 |
| GO:0001520 | outer dense fiber | 5 | 3 | 0.16 | 0.0003 |
| GO:0098826 | endoplasmic reticulum tubular network membrane | 16 | 4 | 0.5 | 0.0013 |
| GO:0030018 | Z disc | 148 | 12 | 4.66 | 0.0025 |
| GO:0032839 | dendrite cytoplasm | 34 | 5 | 1.07 | 0.004 |
| GO:0001725 | stress fiber | 49 | 6 | 1.54 | 0.0042 |
| GO:0034663 | endoplasmic reticulum chaperone complex | 11 | 3 | 0.35 | 0.0042 |
| GO:0031012 | extracellular matrix | 231 | 17 | 7.28 | 0.0054 |
| GO:0032983 | kainate selective glutamate receptor complex | 12 | 3 | 0.38 | 0.0055 |
| GO:0005770 | late endosome | 285 | 18 | 8.98 | 0.0068 |
| GO:0033018 | sarcoplasmic reticulum lumen | 13 | 3 | 0.41 | 0.007 |
| GO:0005859 | muscle myosin complex | 25 | 4 | 0.79 | 0.0073 |
| GO:0005662 | DNA replication factor A complex | 26 | 4 | 0.82 | 0.0084 |
| GO:0009507 | chloroplast | 16 | 3 | 0.5 | 0.0128 |
| GO:0065010 | extracellular membrane-bounded organelle | 6 | 2 | 0.19 | 0.0137 |
| GO:1905103 | integral component of lysosomal membrane | 6 | 2 | 0.2 | 0.0137 |
| GO:0005788 | endoplasmic reticulum lumen | 159 | 13 | 5.0 | 0.0168 |
| GO:0005861 | troponin complex | 7 | 2 | 0.2 | 0.0187 |

Supplementary Table 3 : Top 20 enriched Gene Ontology terms among genes significantly down-regulated in the "fed" condition, compared to the "unfed" condition, for synganglion libraries.

| GO.ID | Term | Annotated | Significant | Expected | weight01 |
|---|---|---|---|---|---|
| **Molecular Function** | | | | | |
| GO:0001227 | DNA-binding transcription repressor activity, RNA polymerase II-s | 127 | 29 | 6.77 | 1.9E-11 |
| GO:0000978 | RNA polymerase II proximal promoter sequence-specific DNA bin | 257 | 33 | 13.7 | 0.0000037 |
| GO:0005089 | Rho guanyl-nucleotide exchange factor activity | 79 | 16 | 4.21 | 0.0000064 |
| GO:0005096 | GTPase activator activity | 246 | 29 | 13.12 | 0.000053 |
| GO:0004971 | AMPA glutamate receptor activity | 11 | 5 | 0.59 | 0.00015 |
| GO:0000977 | RNA polymerase II regulatory region sequence-specific DNA bindi | 435 | 56 | 23.19 | 0.0002 |
| GO:0015485 | cholesterol binding | 58 | 11 | 3.09 | 0.00021 |
| GO:0005516 | calmodulin binding | 217 | 25 | 11.57 | 0.00024 |
| GO:0004143 | diacylglycerol kinase activity | 12 | 5 | 0.64 | 0.00025 |
| GO:1990829 | C-rich single-stranded DNA binding | 7 | 4 | 0.37 | 0.00025 |
| GO:0005244 | voltage-gated ion channel activity | 110 | 23 | 5.86 | 0.00039 |
| GO:0060072 | large conductance calcium-activated potassium channel activity | 8 | 4 | 0.43 | 0.00047 |
| GO:0005001 | transmembrane receptor protein tyrosine phosphatase activity | 21 | 6 | 1.12 | 0.00061 |
| GO:0004970 | ionotropic glutamate receptor activity | 31 | 9 | 1.65 | 0.0008 |
| GO:0005154 | epidermal growth factor receptor binding | 23 | 6 | 1.23 | 0.00104 |
| GO:0020037 | heme binding | 468 | 41 | 24.95 | 0.00127 |
| GO:0008140 | cAMP response element binding protein binding | 5 | 3 | 0.27 | 0.00139 |
| GO:0005509 | calcium ion binding | 569 | 47 | 30.34 | 0.00198 |
| GO:0050839 | cell adhesion molecule binding | 237 | 21 | 12.64 | 0.00199 |
| GO:0070300 | phosphatidic acid binding | 18 | 5 | 0.96 | 0.00204 |
| **Biological Process** | | | | | |
| GO:0000122 | negative regulation of transcription by RNA polymerase II | 563 | 65 | 30.9 | 0.000000017 |
| GO:0007268 | chemical synaptic transmission | 612 | 87 | 33.59 | 0.0000032 |
| GO:0086012 | membrane depolarization during cardiac muscle cell action potent | 14 | 7 | 0.77 | 0.0000036 |
| GO:0045175 | basal protein localization | 14 | 7 | 0.77 | 0.0000036 |
| GO:0008355 | olfactory learning | 36 | 10 | 1.98 | 0.000004 |
| GO:0035023 | regulation of Rho protein signal transduction | 109 | 21 | 5.98 | 0.0000056 |
| GO:0045475 | locomotor rhythm | 41 | 11 | 2.25 | 0.0000068 |
| GO:0019991 | septate junction assembly | 34 | 10 | 1.87 | 0.0000093 |
| GO:1904862 | inhibitory synapse assembly | 5 | 4 | 0.27 | 0.000043 |
| GO:0048147 | negative regulation of fibroblast proliferation | 26 | 8 | 1.43 | 0.000051 |
| GO:0042058 | regulation of epidermal growth factor receptor signaling pathway | 69 | 13 | 3.79 | 0.000054 |
| GO:2000279 | negative regulation of DNA biosynthetic process | 39 | 10 | 2.14 | 0.000054 |
| GO:0007165 | signal transduction | 3588 | 299 | 196.91 | 0.000079 |
| GO:0007585 | respiratory gaseous exchange | 52 | 12 | 2.85 | 0.000085 |
| GO:0046834 | lipid phosphorylation | 48 | 8 | 2.63 | 0.000098 |
| GO:0010944 | negative regulation of transcription by competitive promoter bindin | 10 | 5 | 0.55 | 0.000098 |
| GO:0060244 | negative regulation of cell proliferation involved in contact inhibitio | 10 | 5 | 0.55 | 0.000098 |
| GO:0034765 | regulation of ion transmembrane transport | 228 | 30 | 12.51 | 0.0001 |
| GO:0001738 | morphogenesis of a polarized epithelium | 137 | 12 | 7.52 | 0.00012 |
| GO:0045944 | positive regulation of transcription by RNA polymerase II | 771 | 69 | 42.31 | 0.00015 |
| **Cellular Component** | | | | | |
| GO:0030054 | cell junction | 1061 | 118 | 59.21 | 0.00000011 |
| GO:0098839 | postsynaptic density membrane | 57 | 14 | 3.18 | 0.0000003 |
| GO:0005886 | plasma membrane | 3677 | 320 | 205.2 | 0.00000044 |
| GO:0099056 | integral component of presynaptic membrane | 37 | 11 | 2.06 | 0.0000035 |
| GO:0016323 | basolateral plasma membrane | 251 | 29 | 14.01 | 0.000037 |
| GO:0016020 | membrane | 7989 | 538 | 445.83 | 0.000079 |
| GO:0045211 | postsynaptic membrane | 281 | 44 | 15.68 | 0.00017 |
| GO:0098978 | glutamatergic synapse | 206 | 25 | 11.5 | 0.00021 |
| GO:0005913 | cell-cell adherens junction | 75 | 12 | 4.19 | 0.00031 |
| GO:0098685 | Schaffer collateral - CA1 synapse | 49 | 10 | 2.73 | 0.00031 |
| GO:0005938 | cell cortex | 411 | 35 | 22.94 | 0.00045 |
| GO:0017146 | NMDA selective glutamate receptor complex | 8 | 4 | 0.45 | 0.00056 |
| GO:0014069 | postsynaptic density | 279 | 39 | 15.57 | 0.00057 |
| GO:0043235 | receptor complex | 205 | 31 | 11.44 | 0.00068 |
| GO:0032587 | ruffle membrane | 64 | 11 | 3.57 | 0.00076 |
| GO:1990454 | L-type voltage-gated calcium channel complex | 9 | 4 | 0.5 | 0.00097 |
| GO:0005604 | basement membrane | 90 | 13 | 5.02 | 0.00142 |
| GO:0043005 | neuron projection | 1224 | 121 | 68.3 | 0.0015 |
| GO:0098684 | photoreceptor ribbon synapse | 10 | 4 | 0.6 | 0.00154 |
| GO:0005614 | interstitial matrix | 5 | 3 | 0.3 | 0.00159 |

Supplementary Figure 1: Genomic structure of the gene GABA-1-Rdl, with three predicted isoforms and their relative expression. A: Reannotation of the gene XP_042145571.1 located on a genomic scaffold of *Ixodes scapularis*, NW_024609839.1. The gene (drawing not to scale) has a span of ~247 kbp, on a scaffold of ~92 Mbp. The whole gene is on the minus frame. Boxes correspond to exons. Grey-filled boxes correspond to exons which we consider incorrect (over-predictions in both cases, based on the conserved sequence of GABA-1-Rdl) and by comparison with the homologous sequence in other tick species. Exons in blue correspond to a predicted triplication of one exon (exon 7a, 7b, 7c), whereas only one exon (7b) was annotated for XP_042145571.1. For each numbered exon, the positions indicate the start and end. Numbers followed by a star correspond to reannotations and differ from the published sequence. The translated sequence of each exon is given. B: Alignment of the translated sequences for the three alternative exons 7, including both the sequences for *I. scapularis* based on our reannotation and the homologous sequences from the *I. ricinus* alternative transcripts identified in the synganglion transcriptome (this study). Two trans-membrane domaines are indicated, and an arrow shows the A->S mutations known to confer resistance to dieldrin. C: Relative expression (y-axis) of the three isoforms of GABA-1-Rdl. Expressions was counted as counts per millions with RSEM, and normalized to evaluate relative expression. In x-axis, different synganglion libraries produced in this study (described in Table 1). In blue, red and yellow, estimated relative expression of exon 7a, 7b and 7c respectively.

Supplementary Figure 2: Genomic organization of Histamine-gated-like sequences in *Ixodes scapularis.* We used the Histamine gated-like sequences obtained from our meta-transcriptome of Ixodes ricinus to search homologous genes in *I. scapularis* (homology inferred from near-identity of protein sequences), and to locate them on the genome. The figure shows the entire scaffold NW_024609883 (109.621.045 bp) from the *I. scapularis* genome, which has homologs to His1 to His12 in *I. ricinus*. The upper scale indicates positions in Mbp. The lower scale represents a focus on a smaller region containing clusters of His-like sequences, with a scale in Kbp. Annotated genes and their orientations are indicated by filled triangles, with below, the accession of the protein sequence in *I. scapularis*, and the name of its homologous sequence in *I. ricinus* (this study). A star indicates that the gene model is probably incorrect: XP_042142979 matches with His1 only over the first four exons, while its remaining sequence appears to represent a chimeric fusion with a totally different gene, XP_040070355 is missing an N-term, XP_040070356 is incomplete and matches only the beginning of His3, whereas XP_040068404 matches only the end of His3 and is in opposite frame of XP_ 040070356 (we interpret this as a likely error of the genome assembly, the two accession probably representing respectively the start and end of the same gene). Open triangles indicate regions where no gene has been annotated in *I. scapularis*, but where we detected high similarities with *I. ricinus* genes (for His4 and His9 respectively). For His13 to His18, homologous regions were detected on different scaffolds.

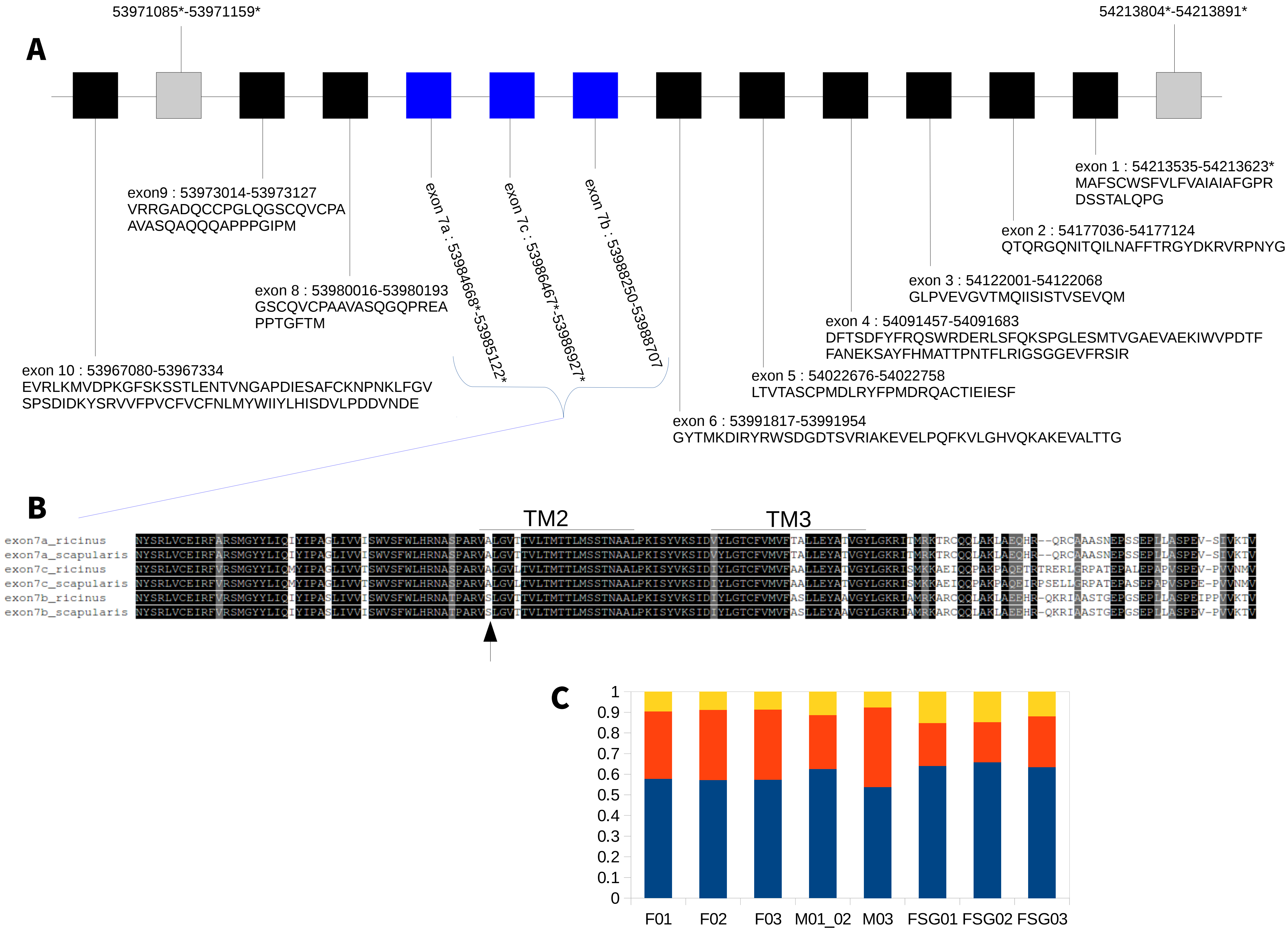
A
53971085*-53971159*
54213804*-54213891*
exon9 : 53973014-53973127
VRRGADQCCPGLQGSCQVCPA
AVASQAQQQAPPPGIPM
exon 8 : 53980016-53980193
GSCQVCPAAVASQGQPREA
PPTGFTM
exon 10 : 53967080-53967334
EVRLKMVDPKGFSKSSTLENTVNGAPDIESAFCKNPNKLFGV
SPSDIDKYSRVVFPVCFVCFNLMYWIIYLHISDVLPDDVNDE
exon 7a : 53984668*-53985122*
exon 7c : 53986467*-53986927*
exon 7b : 53988250-53988707
exon 1 : 54213535-54213623*
MAFSCWSFVLFVAIAIAFGPR
DSSTALQPG
exon 2 : 54177036-54177124
QTQRGQNITQILNAFFTRGYDKRVRPNYG
exon 3 : 54122001-54122068
GLPVEVGVTMQIISISTVSEVQM
exon 4 : 54091457-54091683
DFTSDFYFRQSWRDERLSFQKSPGLESMTVGAEVAEKIWVPDTF
FANEKSAYFHMATTPNTFLRIGSGGEVFRSIR
exon 5 : 54022676-54022758
LTVTASCPMDLRYFPMDRQACTIEIESF
exon 6 : 53991817-53991954
GYTMKDIRYRWSDGDTSVRIAKEVELPQFKVLGHVQKAKEVALTTG
B
TM2
TM3
exon7a_ricinus
exon7a_scapularis
exon7c_ricinus
exon7c_scapularis
exon7b_ricinus
exon7b_scapularis
C
1
0.9
0.8
0.7
0.6
0.5
0.4
0.3
0.2
0.1
0
F01
F02
F03
M01_02
M03
FSG01
FSG02
FSG03

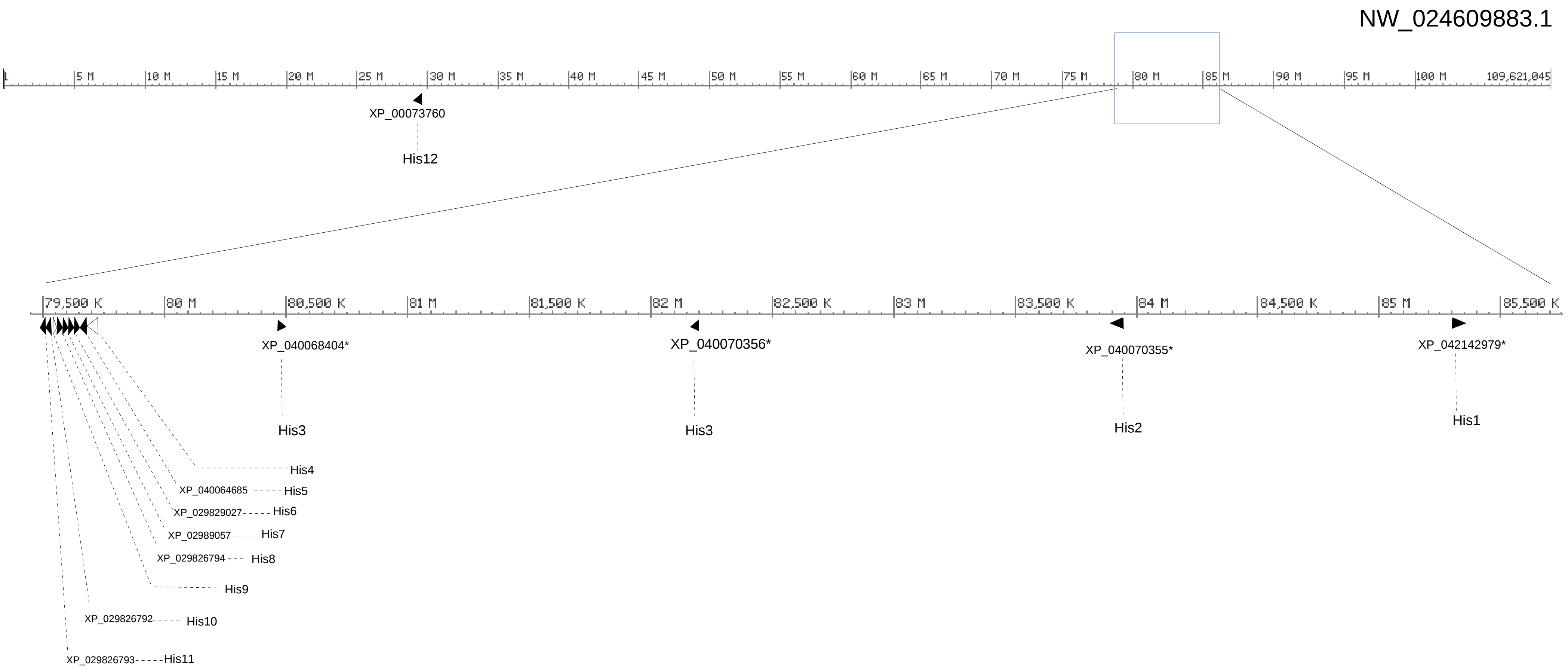

NW_024609883.1
1
5 M
10 M
15 M
20 M
25 M
30 M
35 M
40 M
45 M
50 M
55 M
60 M
65 M
70 M
75 M
80 M
85 M
90 M
95 M
100 M
109,621,045
XP_00073760
His12
79,500 K
80 M
80,500 K
81 M
81,500 K
82 M
82,500 K
83 M
83,500 K
84 M
84,500 K
85 M
85,500 K
XP_040068404*
His3
XP_040070356*
His3
XP_040070355*
His2
XP_042142979*
His1
His4
XP_040064685
His5
XP_029829027
His6
XP_02989057
His7
XP_029826794
His8
His9
XP_029826792
His10
XP_029826793
His11
